# Differential vulnerability of CA1 pyramidal neuron cell types in the 5xFAD Alzheimer’s disease mouse model

**DOI:** 10.64898/2026.05.29.728864

**Authors:** Maricarmen Pachicano, Angela Hurtado, Shrey Mehta, Bayla Breningstall, Yiwen Zhou, Michael S. Bienkowski

## Abstract

Selective neuronal vulnerability is a defining feature of Alzheimer’s disease, yet how neurodegeneration unfolds across defined hippocampal CA1 pyramidal neuron populations remains unresolved. Previously, we showed that CA1 pyramidal neurons are organized into molecularly defined laminar cell types that form stable spatial signatures along the longitudinal axis (Pachicano et al., 2025). Here, we investigated how vulnerability manifests across CA1 pyramidal cell types in the 5xFAD mouse model of amyloid pathology. HiPlex single-molecule fluorescence in situ hybridization was used to quantify gene expression at single-cell resolution across hippocampal CA1 subregions across disease progression. Cells were classified into molecularly defined CA1 pyramidal cell types based on marker gene expression, and changes in density, proportion, and molecular state were assessed. Amyloid-associated pathology differentially affected CA1 pyramidal cell types, revealing distinct trajectories of vulnerability and resilience. Specific populations exhibited early and progressive susceptibility, while other populations demonstrated relative preservation across disease timepoints. These effects were consistent across CA1 subregions, indicating that vulnerability follows cell type identity rather than regional anatomy alone. Together, these findings demonstrate that neurodegeneration in Alzheimer’s disease is structured by intrinsic cell type identity and provide a cellular framework for understanding selective vulnerability.

## Main

The distinct neuroanatomical patterns and slow, progressive spread of pathology in Alzheimer’s disease (AD) suggests that specific brain regions and their constituent cell types are differentially vulnerable or resilient to neurodegeneration. The hippocampal CA1 region is a critical site of early pathology, dysfunction, and cell loss in AD ^1–3^, yet the cellular resolution at which CA1 degeneration unfolds remains poorly understood. Although extensive work has documented changes in neuronal activity, connectivity, and function among CA1 pyramidal neurons in AD models, these studies have largely relied on region-level analyses or coarse anatomical distinctions ^4–8^. Consequently, whether disease progression selectively impacts distinct CA1 pyramidal neuron populations at different stages remains unresolved.

Our previous work developing the Hippocampus Gene Expression Atlas (HGEA) has demonstrated that CA1 pyramidal neurons are organized into at least four molecularly defined gene expression layers that vary systematically along the hippocampal longitudinal axis and correspond to distinct transcriptional identities ^9^. Through distinct laminar architecture, these layers define five CA1 subregions – CA1d (dorsal), CA1i (intermediate), CA1v (ventral), CA1vv (ventral-ventral), and CA1c (caudal). This laminar gene expression framework provides a refined approach for resolving pyramidal neuron diversity beyond traditional anatomical classifications such as superficial versus deep layers ^10,11^. In contrast to conventional AD neurodegeneration studies that quantify neuronal loss using Nissl or NeuN-based staining to generate region-wide counts ^12,13^, this approach enables molecularly defined, cell type-specific labeling for quantification. Whether this laminar molecular architecture can identify selective vulnerability in AD remains unknown.

To address these questions, we examined CA1 pyramidal neuron organization in the 5xFAD mouse model of AD, a well-characterized amyloid-driven model carrying five familial AD mutations that exhibits progressive amyloid pathology across aging timepoints ^14,15^. Leveraging the established HGEA CA1 cell type definitions ^9,10^ and RNAscope-based single-cell analysis, we quantified laminar gene expression, cellular composition, and molecular state across disease progression in male and female 5xFAD mice. Sample sizes by age, sex, and HGEA level are summarized in Fig. S1.

Our primary analyses focused on a mid-rostrocaudal level of the CA1 where all four laminar-defined pyramidal neuron populations are present, enabling direct comparison within a shared anatomical context. By integrating laminar gene expression mapping with quantitative cellular and molecular analyses across longitudinal position, disease stage, and sex, this study establishes a cell type-resolved framework for identifying divergent trajectories of vulnerability and resilience among molecularly distinct CA1 pyramidal neuron cell type populations.

## Results

### Conserved, but dysregulated CA1 laminar gene expression organization in 2-month-old WT and 5xFAD CA1

Using previously established CA1 laminar marker genes (*Lrmp*, *Ndst4*, *Trib2*, and *Peg10*), we examined laminar gene expression profiles of CA1 pyramidal neurons in WT (wild-type) and 5xFAD mice at 2 months-old (mo) (Fig. 1). At this early disease stage, the overall laminar gene expression signatures distinguishing CA1 subregions were preserved between both genotypes, despite differences in the magnitude and variability of individual marker gene expression.

**Fig 1.**
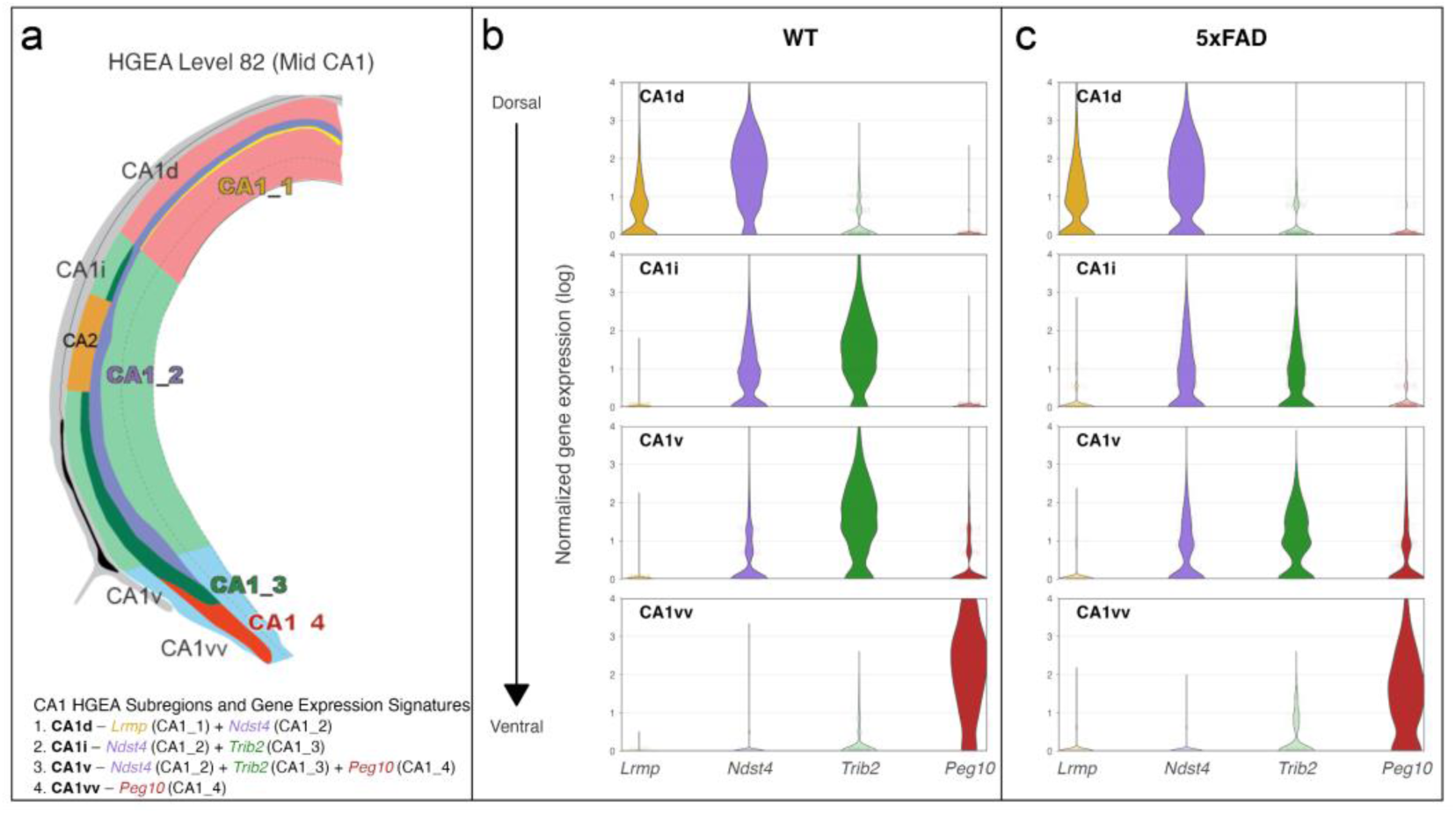
Laminar gene expression signatures are preserved in 2mo WT and 5xFAD mice. **a,** HGEA Level 82 (mid-CA1), illustrating CA1 subregions (CA1d, CA1i, CA1v, CA1vv) along the dorsoventral axis and the spatial organization of laminar gene expression signatures across the pyramidal cell layer, which contains four molecularly defined populations (CA1_1–CA1_4). Color-coded regions indicate characteristic laminar gene expression patterns for CA1_1 marker gene *Lrmp* (yellow), CA1_2 marker gene *Ndst4* (purple), CA1_3 marker gene *Trib2* (green), and CA1_4 marker gene *Peg10* (red), which together define a laminar cell type-resolved molecular framework. **b,** Violin plots of log-normalized gene expression of *Lrmp*, *Ndst4*, *Trib2*, and *Peg10* across CA1 subregions in 2mo WT mice. **c,** Violin plots showing log-normalized gene expression of the same genes across CA1 subregions in 2mo 5xFAD mice. Gene-subregion combinations corresponding to expected laminar expression patterns are shown at full opacity, whereas non-corresponding combinations are displayed with reduced opacity for visual context. Violin plots represent single-cell measurements from WT (n=4 mice) and 5xFAD (n=5 mice) animals. Source data are provided with this figure.

At HGEA coronal level 82, all four CA1 laminar gene expression layers are differentially distributed along the longitudinal axis to define each CA1 subregion’s unique gene expression profile. Specifically, CA1d is a bilaminar region defined by enriched *Lrmp* and *Ndst4* expression, CA1i is also a bilaminar region but characterized by *Ndst4* and *Trib2* expression, CA1v is trilaminar exhibiting complementary layers of *Ndst4*, *Trib2*, and *Peg10* expression, and CA1vv is a monolayer dominated by *Peg10* expression (Fig. 1a). Gene expression profiles of these CA1 subregions were highly similar between 2-month-old WT and 5xFAD mice (Fig. 1b,c) although modest differences in expression magnitude were evident. Notably, *Lrmp* appears modestly upregulated in CA1d, whereas Trib2 expression appears reduced in CA1i and CA1v in 5xFAD mice relative to WT. In contrast, *Ndst4* expression distributions appear broadly similar between genotypes. This preservation of laminar gene expression signatures at the early 2mo timepoint establishes a shared molecular baseline between WT and 5xFAD mice, while suggesting that CA1 neurons in 5xFAD mice already exhibit dysregulated gene expression prior to the development of extracellular amyloid plaques.

### Laminar gene expression organization across disease progression in 5xFAD mice

Using topographic maps of CA1 pyramidal cells at mid-rostrocaudal sections (HGEA Level 82), we visualized laminar gene expression organization in 5xFAD mice at 2, 6, and 14mo (Fig. 2a-c). These maps depict the spatial distribution of CA1 laminar marker gene expression across dorsoventral and mediolateral axes and provide a descriptive overview of laminar gene expression organization across disease progression. At 2mo (Fig. 2a), laminar gene expression patterns were spatially organized in a manner consistent with the revised HGEA ^9^. The overall spatial organization of laminar gene expression layers was maintained at 6 and 14mo (Fig. 2b,c), although changes in gene expression density occurred with disease progression.

**Fig 2.**
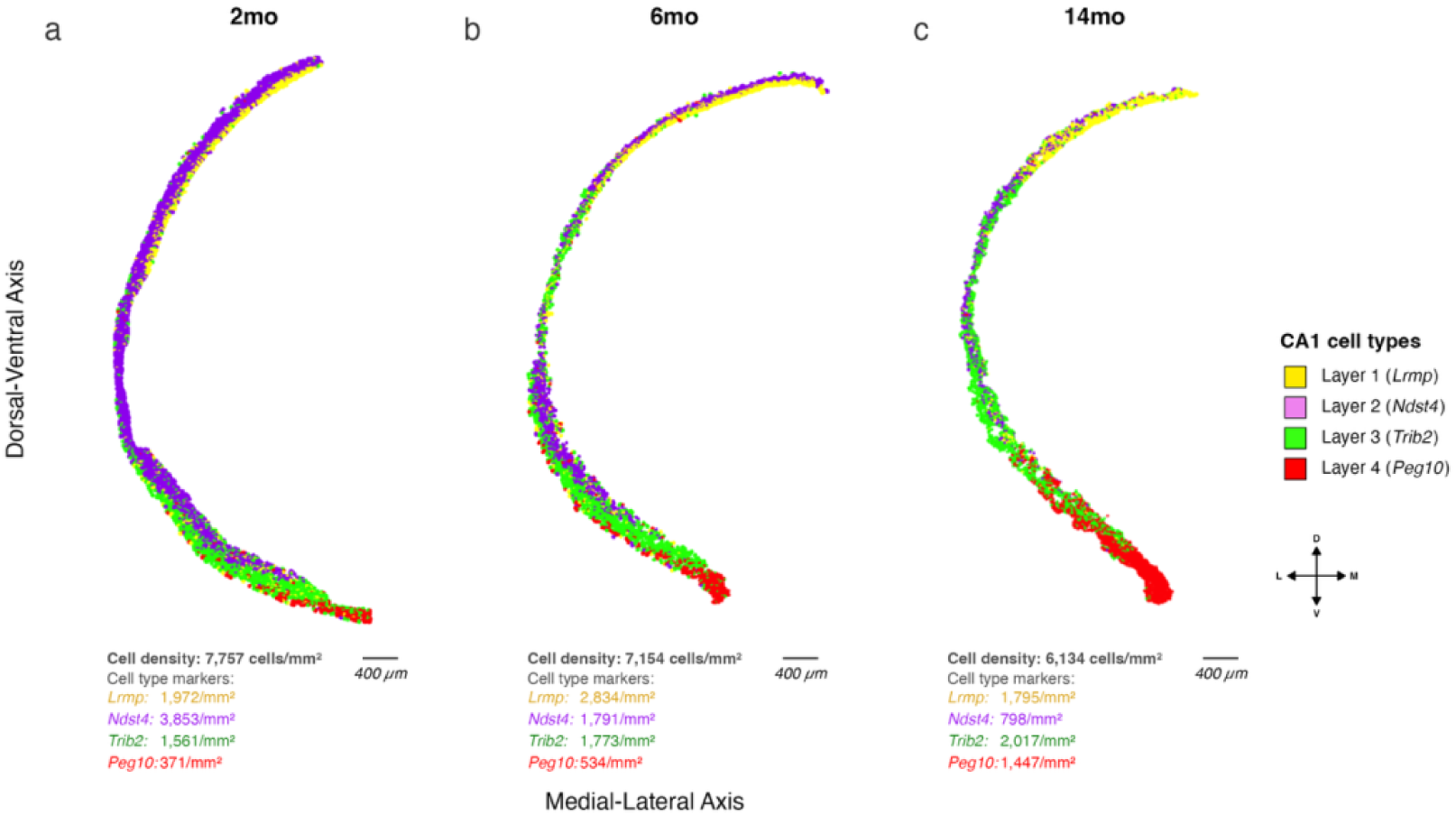
Spatial topography of CA1 laminar gene expression patterns across disease progression in 5xFAD mice. CA1 laminar topography maps from male 5xFAD mice aged 2mo **(a)**, 6mo **(b)**, and 14mo **(c)**, at the mid-rostrocaudal level (HGEA Level 82). Individual cells are plotted according to their mediolateral (x-axis) and dorsoventral (y-axis) anatomical positions within the CA1 pyramidal layer. Crosshairs denote anatomical axis orientation (D, dorsal; V, ventral; M, medial; L, lateral). Cells are colored according to the gene with the highest log-normalized expression value, corresponding to laminar cell type populations associated with *Lrmp* (Layer 1, yellow), *Ndst4* (Layer 2, purple), *Trib2* (Layer 3, green), and *Peg10* (Layer 4, red). Cells lacking detectable expression across markers are unassigned. Total cell density and marker gene-associated cell densities (cells/mm^2^) are indicated below each map. Scale bars, 400 μm. Source data are provided with this figure.

Consistent with these topographic representations, RNAscope single molecule fluorescent *in situ hybridization* (smFISH) confirmed clear laminar segregation of CA1 marker gene RNA transcript expression at single-cell resolution across all ages examined (Fig. 3a-c). High-resolution images demonstrated reproducible spatial positioning of *Lrmp*, *Ndst4*, *Trib2*, and *Peg10* expression within the pyramidal layer across CA1 subregions, with each marker occupying distinct laminar positions at 2, 6, and 14mo (Fig. 3d-g). Collectively, the concordance between topographic gene expression maps and RNAscope spatial signal confirms that the foundational laminar organization of CA1 is preserved across disease progression at the mid-CA1 level.

**Fig 3.**
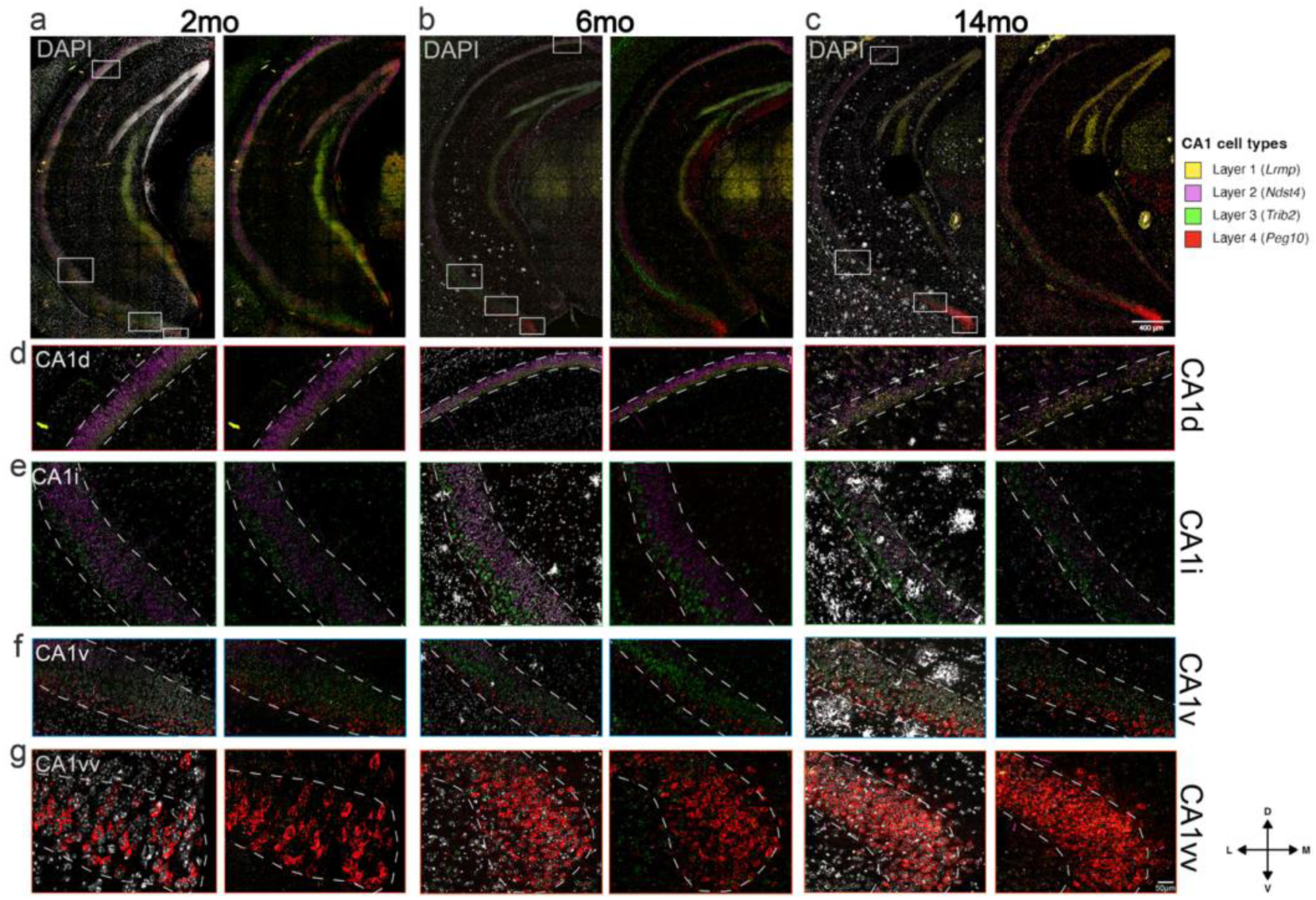
RNAscope imaging of CA1 laminar gene expression across disease progression in male 5xFAD mice. 2mo **(a)**, 6mo **(b)**, and 14mo **(c)**, corresponding to HGEA Level 82. DAPI (grayscale) shows cytoarchitecture, color channels show RNAscope signal for laminar marker genes. **(d-g)** High-magnification views of CA1 subregions CA1d **(d)**, CA1i **(e)**, CA1v **(f)**, and CA1vv **(g)**, corresponding to the boxed regions in panels **(a-c)**. Each row shows the same subregion across all three ages (2mo, 6mo, 14mo). White dashed lines delineate the boundaries of the CA1 pyramidal layer. Crosshairs indicate dorsoventral and mediolateral orientation of the images (D, dorsal; V, ventral; M, medial; L, lateral). Scale bars, 400 μm **(a-c)** and 50 μm **(d-g)**.

### Rostrocaudal variation in CA1 laminar gene expression organization

To further evaluate the robustness of the CA1 architecture, we examined laminar organization along the rostrocaudal axis by generating topographic maps at rostral and caudal levels across disease progression (2mo, 6mo, and 14mo) in 5xFAD mice (Fig. S2). These maps extend the mid-CA1 laminar characterization to additional rostrocaudal levels, visualizing the changes to the spatial arrangement of laminar gene expression layers along the longitudinal axis. At rostral CA1 levels, where laminar gene expression is dominated by *Lrmp* and *Ndst4*, topographic maps revealed relatively stable spatial organization of superficial and deep gene expression layers across disease progression (Fig. S2a). The relative positioning and continuity of these laminar gene expression sublayers were maintained across ages, indicating that rostral CA1 laminar organization is largely preserved despite disease progression.

By contrast, more caudal CA1 regions exhibited greater disease-associated variability. At the mid-rostrocaudal CA1 level (HGEA Level 82), where all four gene expression layers are present, the overall spatial arrangement of these domains remained apparent across ages, though shifts in visual prominence of specific gene expression layers became evident with disease progression (Fig. 2). These findings indicate differential rostrocaudal disruption of laminar organization within the 5xFAD model. This spatial specificity may explain why prior studies that examined a single coronal level did not report level-dependent differences in CA1 vulnerability ^16,17^.

At caudal CA1 (HGEA Level 89), where CA1 adopts a trilaminar gene expression organization composed of *Trib2*, *Ndst4*, and *Lrmp* gene expression layers (deep to superficial), laminar gene expression structure was similarly retained across disease progression, despite a progressive decline in overall cellular representation within the pyramidal layer (Fig. S2b). Changes were most evident within *Ndst4*-expressing layers, whereas *Lrmp*-associated superficial layers were comparatively preserved. Together, these analyses reveal spatial heterogeneity in laminar vulnerability along the rostrocaudal axis, highlighting regions of relative preservation versus disruption across disease progression.

### Disease-associated changes in CA1 laminar gene expression

To assess how disease-associated changes affect marker gene expression, we analyzed pooled single-cell gene expression profiles across CA1 subregions in 2, 6, and 14mo 5xFAD mice (Fig. 4a). Expression of *Lrmp*, primarily localized in Layer 1 cells of CA1d, increased modestly with disease progression. In contrast, the layer 2 cell marker *Ndst4* substantially declined over disease timepoints in CA1d, CA1i, and CA1v. *Trib2* expression also progressively declined within CA1i and CA1v whereas *Peg10* expression remained stable or increased within CA1v and CA1vv. Direct comparison of 2mo and 14mo stages highlighted these profile-level changes (Fig. 4b). Split-violin analyses revealed pronounced reductions in *Ndst4* and *Trib2* expression across CA1 subregions, alongside increases in *Lrmp* and *Peg10* expression. Together, these analyses demonstrate the shifting landscape of CA1 gene expression over the disease time course, presenting challenges to consistent cell type identification amid varying levels of marker genes and their coexpression within individual cells.

**Fig. 4.**
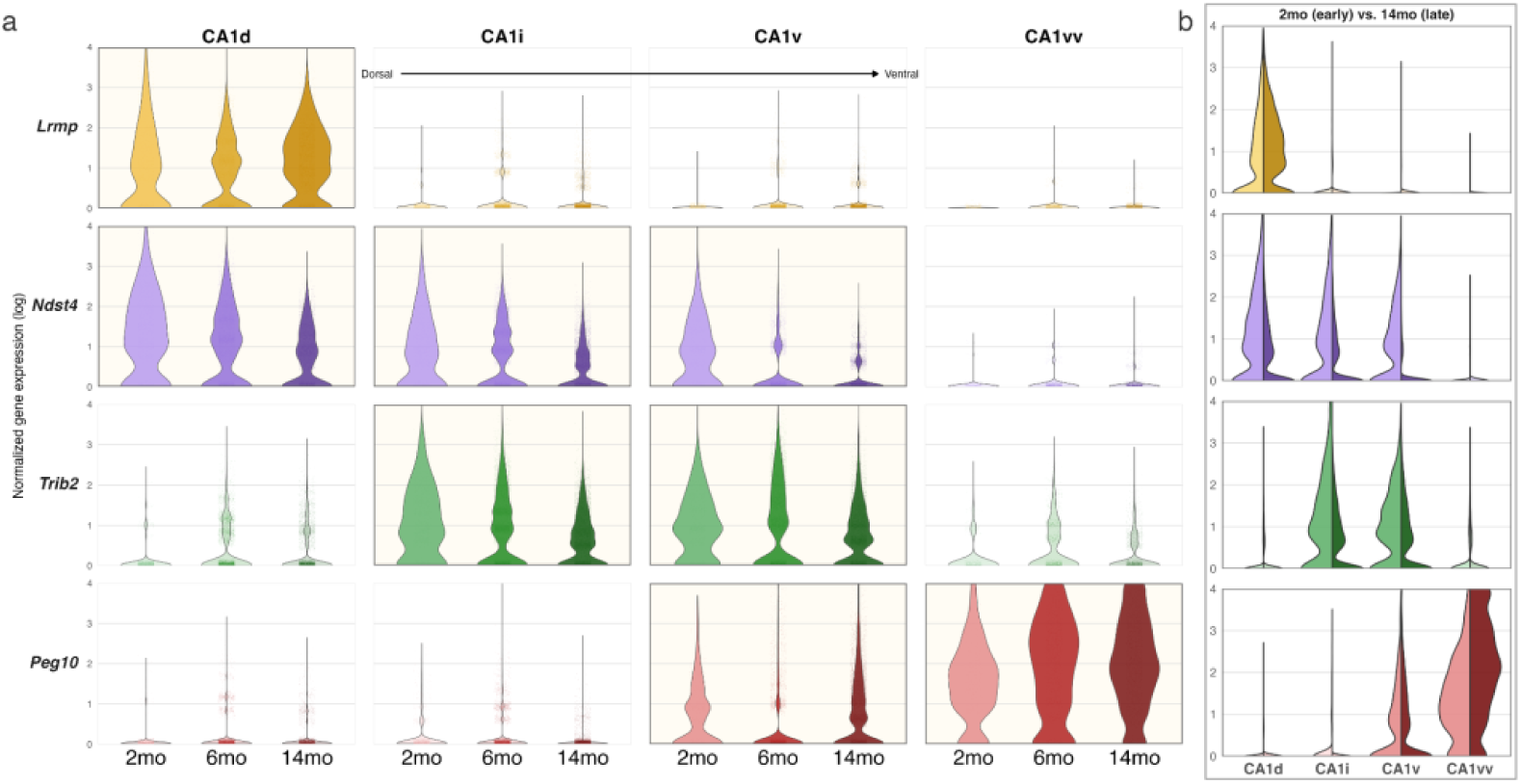
Gene expression distributions across CA1 subregions and ages in 5xFAD mice. **a,** Violin plots showing the distribution of log-normalized gene expression for *Lrmp*, *Ndst4*, *Trib2*, and *Peg10* (rows) across dorsal-to-ventral CA1 subregions (CA1d, CA1i, CA1v, CA1vv) in 5xFAD mice at 2mo, 6mo, and 14mo. Expression distributions are shown separately for each gene and subregion, with age groups displayed along the x-axis. Marker gene-subregion combinations corresponding to expected laminar gene expression patterns are shown at full opacity, while non-corresponding combinations are displayed with reduced opacity for visual context. Cells from all animals within each age group were pooled; sample sizes were 2mo, n=5 (2 male, 3 female); 6mo, n=6 (3 male, 3 female); 14mo, n=9 (4 male, 5 female). **b,** Split violin plots comparing log-normalized gene expression between early (2mo) and late (14mo) 5xFAD mice for each CA1 subregion. Each split violin represents the distribution of per-cell expression values for the indicated gene within a given subregion. Source data are provided with this figure.

Given the pronounced gene expression differences observed at advanced disease stage, we next examined whether laminar marker genes exhibit coordinated expression within individual CA1 pyramidal neurons at 14mo using a SCAMPR-based coexpression framework (Fig. S3), building on the structured laminar coexpression relationships previously defined in the wild-type CA1 pyramidal layer (Pachicano et al., 2025, Fig. S5). Pairwise analyses revealed coordinated and spatially structured relationships between neighboring laminar markers across CA1 subregions, with continuous trajectories in coexpression space that were maintained despite disease-associated shifts in expression. These coordinated patterns indicate that laminar profiles reflect organized molecular states rather than independent gene-level changes. However, these analyses do not distinguish whether reduced marker expression reflects loss of specific pyramidal neuron populations or transcriptional downregulation within surviving cells. To resolve this distinction, spatial analyses must be performed at the level of defined CA1 pyramidal neuron cell types rather than gene expression alone.

### Laminar framework for analysis of cell type-specific vulnerability and resilience

To transition from gene expression distributions to cell type-resolved analyses, the CA1 HGEA was used as a laminar framework for defining pyramidal neuron cell types in the 5xFAD mouse model (Fig. 5a). This framework provides a spatial reference for assigning individual pyramidal neurons to laminar-defined cell types based on marker expression.

**Fig. 5.**
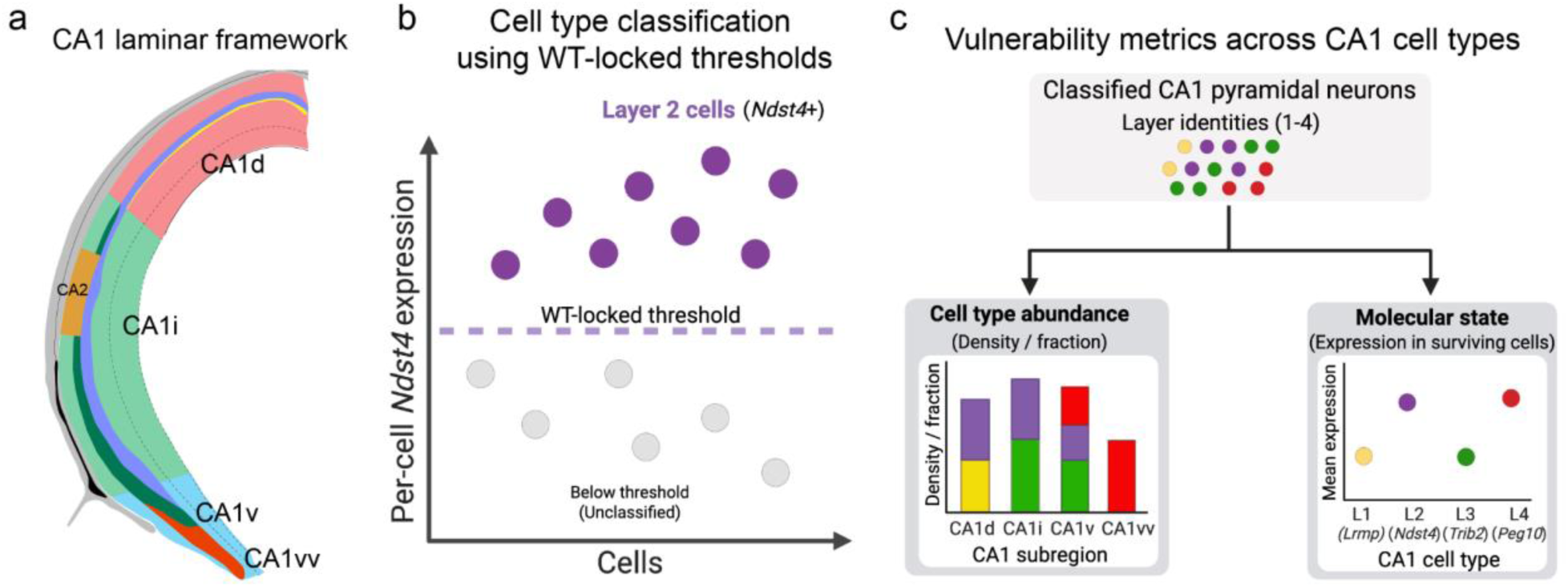
Framework for cell type-resolved analysis of CA1 vulnerability. **a,** HGEA Level 82 showing the CA1 laminar framework across subregions (CA1d, CA1i, CA1v, and CA1vv) along the dorsoventral axis and the spatial organization of gene-defined pyramidal neuron layers. **b,** Conceptual schematic illustrating classification of individual cells into laminar CA1 pyramidal neuron types using WT-anchored gene expression thresholds, shown here for the Layer 2 marker *Ndst4* as an example. Cells exceeding the WT-derived threshold for a given marker are classified as belonging to the corresponding cell type, with cells exceeding thresholds for multiple markers assigned to the dominant laminar marker. *Created in BioRender. Pachicano, M. (2026)* https://BioRender.com/6pexoq8. **c,** Conceptual representation of vulnerability metrics applied to classified CA1 pyramidal neuron populations. Cell type abundance is assessed as the density or fraction of marker-positive cells across CA1 subregions, while molecular state reflects the mean gene expression within surviving cells of each classified laminar population. *Created in BioRender. Pachicano, M. (2026)* https://BioRender.com/l8yjgw5.

Using per-cell marker expression measurements, pyramidal neurons were classified according to the established CA1 layer-subregion correspondence defined by layer-enriched markers (*Lrmp, Ndst4, Trib2, Peg10*) (Fig. 5b). Gene-specific positivity thresholds derived from pooled 2mo WT reference animals were applied uniformly across all disease stages to assign cells to laminar identities (See methods), anchoring classification to a stable WT-defined reference rather than to disease-associated shifts in expression.

This framework enabled systematic, cell type-resolved quantification of vulnerability across disease stages using two complementary readouts: (i) cell type abundance, measured as the density and fraction of positive cells, and (ii) molecular state, assessed as mean normalized gene expression within surviving cells (Fig. 5c). Together, these metrics distinguish changes in cellular representation from changes in transcriptional state within defined CA1 pyramidal neuron populations.

### Disease-associated changes in CA1 laminar cell type density

Using the laminar classification framework described above, we quantified positive-cell densities of CA1 pyramidal neuron cell types across CA1 subregions and disease stages at mid-CA1 (HGEA Level 82) (Fig. 6). Disease-stage effects were evaluated using Kruskal–Wallis tests with Benjamini–Hochberg FDR correction across corresponding layer–subregion panels. Parallel analyses of cell type fraction and molecular state across all CA1 pyramidal neuron cell types and subregions are shown in Fig. S4.

**Fig 6.**
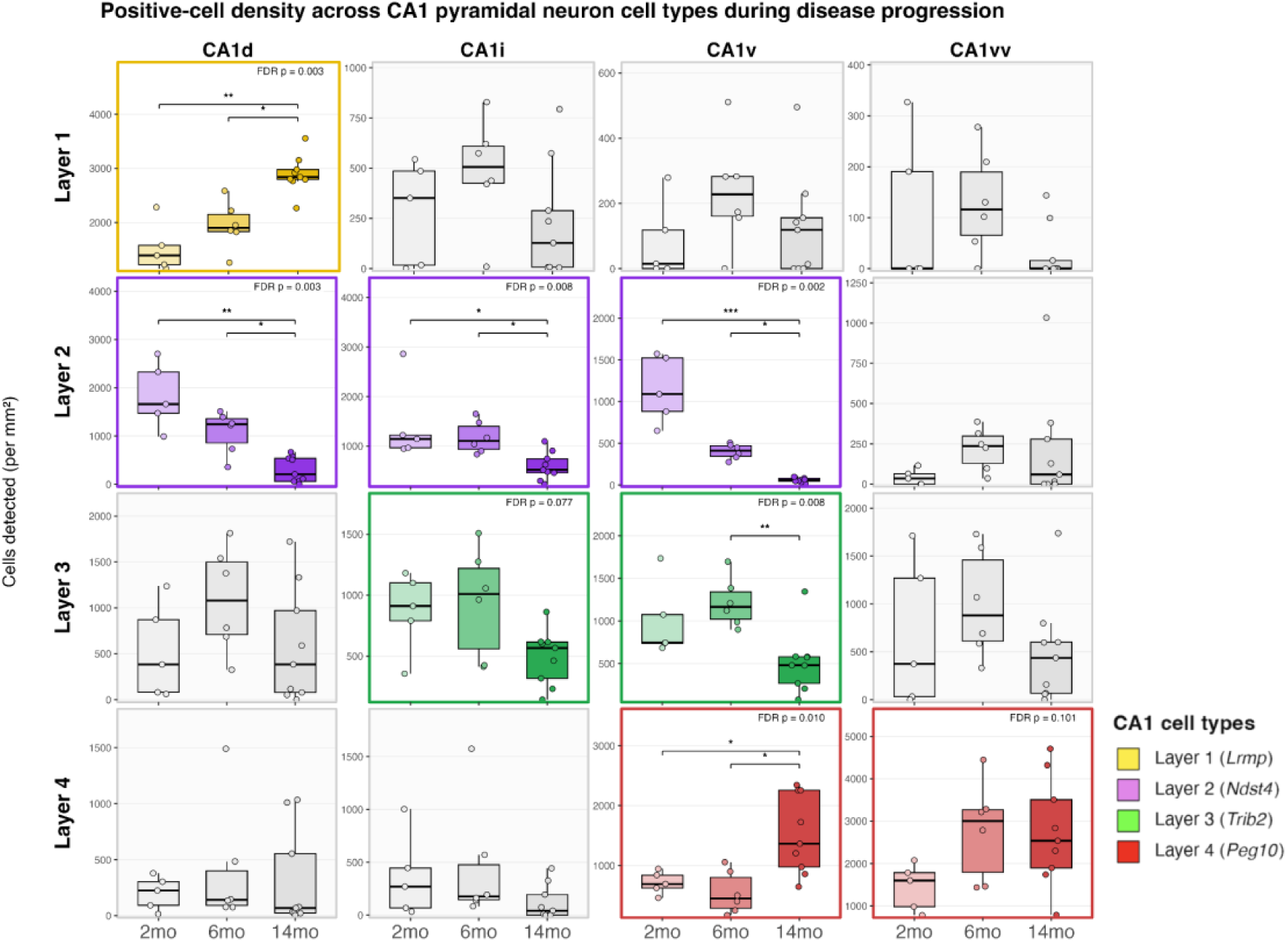
Cell type- and subregion-specific changes in CA1 laminar-defined cell type densities across disease progression in 5xFAD mice. Boxplots summarize per-animal positive-cell densities, with individual points representing single animals. Quantification is based on HGEA Level 82 (mid-CA1) sections with the following sample sizes: 2mo, n=5 (2 male, 3 female); 6mo, n=6 (3 male, 3 female); 14mo, n=9 (4 male, 5 female). Positive cells were defined using gene-specific expression thresholds derived from pooled 2mo WT reference animals. Layer-subregion combinations corresponding to the expected laminar cell type distribution are highlighted with colored borders and gene-tinted fills (lighter to darker with disease stage), whereas non-corresponding gene-subregion combinations are shown in grayscale. Statistical significance was assessed using Kruskal-Wallis tests comparing 2mo, 6mo, and 14mo 5xFAD mice within each corresponding cell type–subregion panel, with Benjamini-Hochberg FDR correction applied across tested panels. Where significant effects were detected, pairwise comparisons were performed using Dunn’s test. FDR-adjusted p-values are shown, and significant pairwise comparisons are indicated by brackets and asterisks. Exact p-values and pairwise Dunn post-hoc comparisons are provided in the corresponding Source Data file.

Because each marker labels a subset of CA1 pyramidal neurons, the proportion of cells exceeding the WT-anchored threshold varied by gene and subregion. Gene-specific thresholds derived from pooled 2mo WT reference animals were held constant across ages to allow direct comparison of laminar cell type identity across disease stages, such that cells falling below the threshold at later ages reflect either neuronal loss or reduced expression of the defining marker. Within the subregions containing the corresponding laminar cell type, approximately 40-60% of cells exceeded the threshold across animals and ages, whereas <10% of cells exceeded threshold in subregions lacking that cell type, consistent with the spatial restriction of CA1 laminar organization. Spatial maps of marker gene expression across CA1 subregions and disease stages are shown in Fig. S5.

CA1 Layer 1 (CA1_1) cell types, restricted to the CA1d at this coronal level, showed a significant increase in positive-cell density with disease progression. In contrast, CA1 Layer 2 (CA1_2) cell types, which span CA1d, CA1i, and CA1v, showed significant reductions in positive-cell density across subregions, with the most pronounced decreases observed in CA1d. CA1 Layer 3 (CA1_3) cell types showed a more limited pattern of change, with a significant disease-stage effect detected in CA1v but not CA1i (although this effect may be trending). CA1 Layer 4 (CA1_4) cell types showed increased positive-cell density in CA1v, whereas changes in CA1vv did not reach significance. Dunn’s post hoc test indicated that, in nearly all significant panels, disease-stage effects were driven by differences involving the 14mo group, whereas 2mo and 6mo animals generally did not differ significantly. These analyses demonstrate differential vulnerability of CA1 pyramidal neuron cell types in AD, with CA1 Layer 2 pyramidal neurons representing the most consistently vulnerable population, while CA1 Layers 1 and 4 exhibit relative preservation.

### Susceptibility of CA1 Layer 2 pyramidal neurons

To determine whether specific CA1 pyramidal neuron populations exhibit differential vulnerability to disease progression, we next examined cellular and molecular changes within the CA1 Layer 2 population across subregions and disease timepoints at mid-CA1 (HGEA Level 82). CA1 Layer 2 pyramidal neurons emerged as the most consistently affected population, exhibiting layer- and subregion-specific reductions across disease progression.

Quantification of cell type fractions revealed robust disease-associated reductions in the proportion of CA1 Layer 2 cells in CA1d, CA1i, and CA1v, with the largest decreases observed in dorsal and intermediate CA1 subregions (Fig. 7a). Dunn’s post hoc test indicated that reductions were primarily driven by differences involving the 14mo group, whereas 2mo and 6mo animals generally did not differ significantly. Reductions in Layer 2 cell fraction were accompanied by parallel declines in *Ndst4* expression within classified cells (Fig. 7b), indicating that both transcriptional downregulation and loss of classified cells contribute to the observed vulnerability.

**Fig 7.**
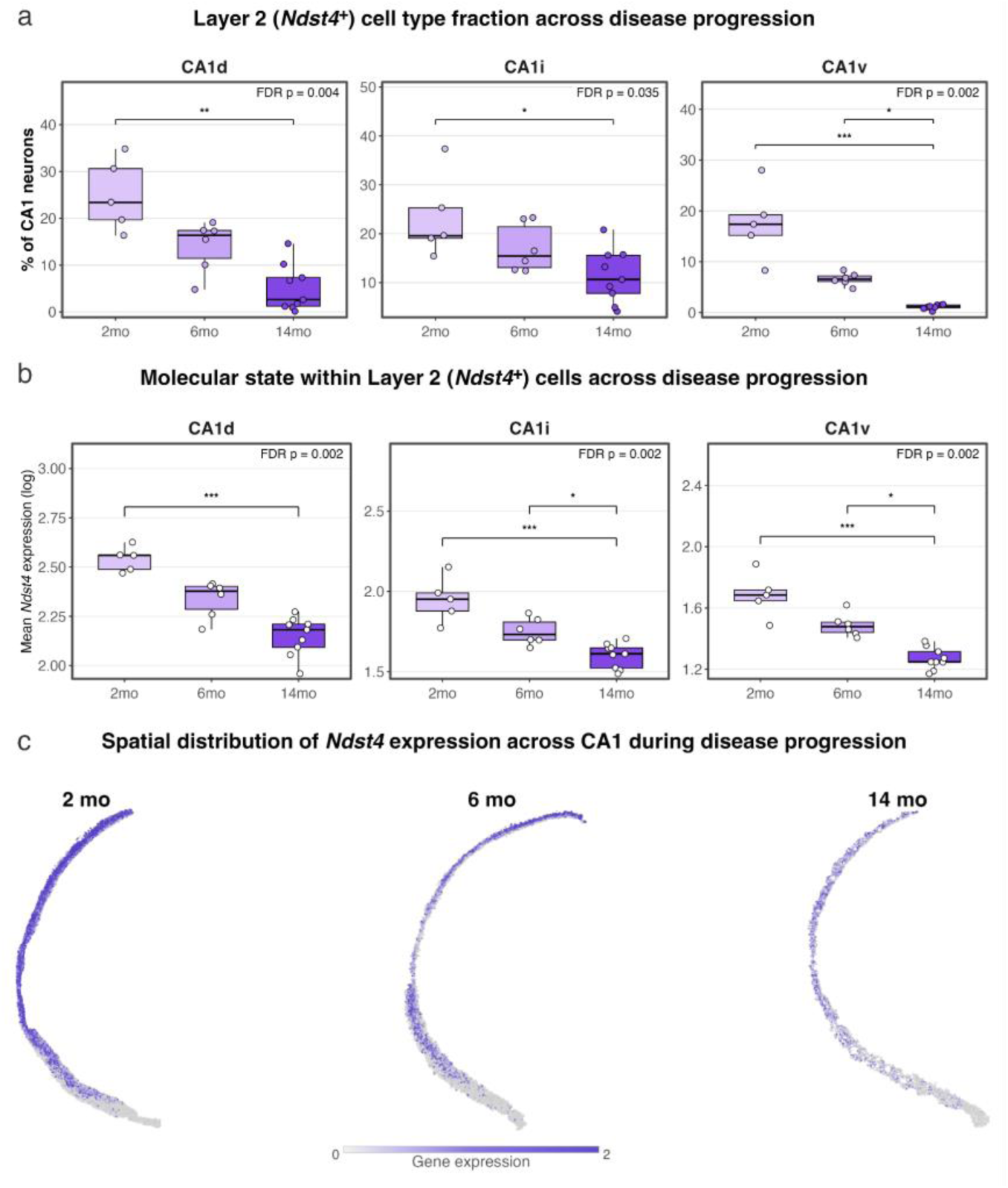
Disease-associated susceptibility of CA1 Layer 2 pyramidal neurons in 5xFAD mice. **a,** Fraction of the CA1 Layer 2 pyramidal neurons, expressed as the percentage of total CA1 pyramidal neurons within subregions where this layer is detected (CA1d, CA1i, CA1v), across timepoints (2mo, 6mo, 14mo) in 5xFAD mice. **b,** Molecular state of CA1 Layer 2 pyramidal neurons, quantified as mean *Ndst4* expression within classified Layer 2 cells across the same subregions and timepoints. For panels **a** and **b**, boxplots summarize per-animal values, with individual points representing single animals. Quantification is based on HGEA Level 82 (mid-CA1) sections with the following sample sizes: 2mo, n=5 (2 male, 3 female); 6mo, n=6 (3 male, 3 female); 14mo, n=9 (4 male, 5 female). **c,** Spatial maps showing the distribution of *Ndst4* gene expression across CA1 at 2mo, 6mo, and 14mo 5xFAD mice, displayed within the same laminar framework. These maps visualize gene expression across all pyramidal neurons and provide anatomical context for the quantitative analyses in panels **a** and **b**. Statistical significance for panels **a** and **b** was assessed using Kruskal-Wallis tests comparing 2mo, 6mo, and 14mo 5xFAD mice within each corresponding cell type–subregion panel, with Benjamini-Hochberg FDR correction applied across tested panels. Where significant effects were detected, pairwise comparisons were performed using Dunn’s test. FDR-adjusted p-values are shown, and significant pairwise comparisons are indicated by brackets and asterisks. Exact p-values and pairwise Dunn post-hoc comparisons are provided in the corresponding Source Data file.

To distinguish disease-associated changes from those attributable to normative aging, age-matched littermate WT animals were analyzed at the aged endpoint (14mo) using the same classification framework (Fig. S6). In contrast to the pronounced, layer-selective reductions observed in 5xFAD mice, age-matched WT animals exhibited comparatively modest and less laminar-specific changes in cell type density, fraction, and molecular state, indicating that the laminar vulnerability pattern is primarily associated with disease rather than aging alone.

Spatial maps confirmed progressive contraction of *Ndst4* signal intensity and spatial coverage with advancing disease (Fig. 7c), providing anatomical context for the observed reductions in Layer 2 cell fraction and molecular state. Finally, coexpression analyses described above supported coordinated molecular disruption within vulnerable CA1 pyramidal neuron populations, as relationships between laminar marker genes were altered in regions exhibiting reduced Layer 2 representation at 14mo (Fig. S3c). Together, these convergent cellular, molecular, and spatial analyses identify CA1 Layer 2 pyramidal neurons as a selectively susceptible cell type during disease progression in the 5xFAD mouse model.

### Preservation of CA1 Layer 1 and Layer 4 pyramidal neurons

In contrast to CA1 Layer 2, Layer 1 and Layer 4 pyramidal neurons exhibited stable or increased cellular representation across disease progression in 5xFAD mice (Fig. 8a). In dorsal CA1 (CA1d), the proportion of CA1 Layer 1 (*Lrmp*^+^) cells increased significantly with disease progression, with the largest shift observed at 14mo. This change in cellular composition was accompanied by a significant increase in mean *Lrmp* expression within classified Layer 1 cells (Fig. 8b). Consistent with these quantitative findings, spatial maps of *Lrmp* expression illustrated an expanded distribution of signal along the superficial CA1 pyramidal layer at later disease stages (Fig. 8c).

**Fig 8.**
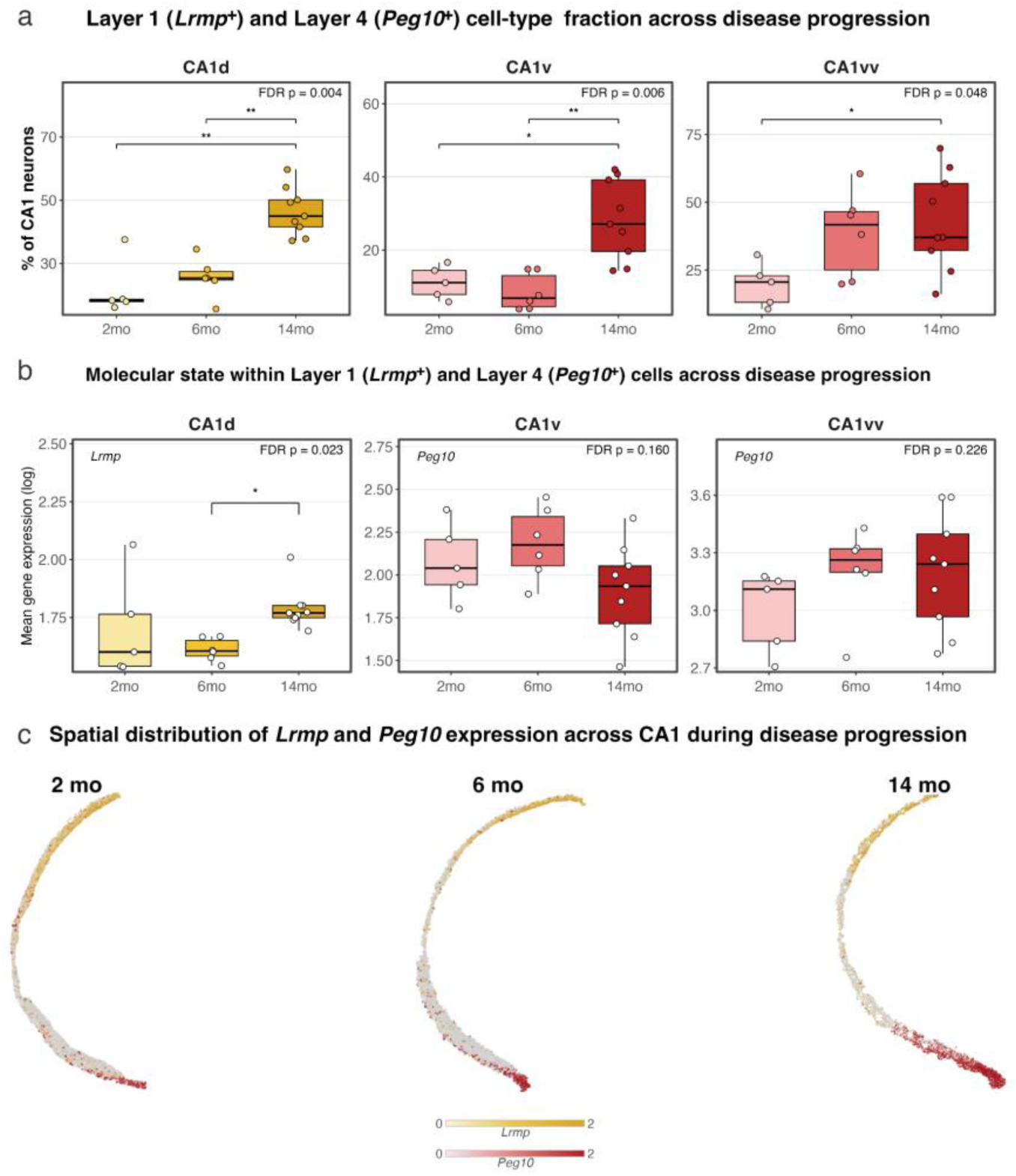
Disease-associated resilience of CA1 Layer 1 and Layer 4 pyramidal neurons in 5xFAD mice. **a,** Fraction of CA1 Layer 1 and Layer 4 pyramidal neurons across timepoints (2mo 6mo, and 14mo) in 5xFAD mice. Values are expressed as the percentage of total CA1 pyramidal neurons within subregions where these layers are detected (CA1d for Layer 1; CA1v and CA1vv for Layer 4). **b,** Molecular state of CA1 Layer 1 and Layer 4 pyramidal neurons, quantified as mean log-normalized *Lrmp* or *Peg10* expression within classified Layer 1 or Layer 4 cells across the same subregions and timepoints. For panels **a** and **b**, boxplots summarize per-animal values, with individual points representing single animals. Quantification is based on HGEA Level 82 (mid-CA1) sections with the following sample sizes: 2mo, n=5 (2 male, 3 female); 6mo, n=6 (3 male, 3 female); 14mo, n=9 (4 male, 5 female). **c,** Spatial maps showing the distribution of *Lrmp* and *Peg10* expression across CA1 at 2mo, 6mo, and 14mo in 5xFAD mice, displayed using the same laminar framework. These maps visualize gene expression across all pyramidal neurons and provide anatomical context for the quantitative analyses in panels **a** and **b**. Statistical significance for panels **a** and **b** was assessed using Kruskal-Wallis tests comparing 2mo, 6mo, and 14mo 5xFAD mice within each corresponding cell type–subregion panel, with Benjamini-Hochberg FDR correction applied across tested panels. Where significant effects were detected, pairwise comparisons were performed using Dunn’s test. FDR-adjusted p-values are shown, and significant pairwise comparisons are indicated by brackets and asterisks. Exact p-values and pairwise Dunn post-hoc comparisons are provided in the corresponding Source Data file.

CA1 Layer 4 pyramidal neurons similarly exhibited significant increases in relative abundance over disease progression within ventral CA1 subregions, including CA1v and CA1vv (Fig. 8a). In contrast to the changes in cellular representation, mean *Peg10* expression within classified Layer 4 cells remained relatively stable across disease stages (Fig. 8b). Spatial maps of *Peg10* expression revealed modest expansion of signal within ventral CA1 at later stages (Fig. 8c). Dunn’s post hoc test indicated that significant differences across these panels were primarily driven by contrasts involving the 14mo disease stage, whereas 2mo and 6mo animals did not differ significantly. These analyses indicate that CA1 Layer 1 and Layer 4 pyramidal neuron populations maintain – or expand – their cellular representation during disease progression in 5xFAD mice, in contrast to the pronounced vulnerability observed in CA1 Layer 2.

### Sex-specific differences in CA1 laminar responses emerge at advanced disease stage

To assess whether sex differences influence the vulnerability of CA1 pyramidal neurons during disease progression, we compared male and female topographic maps of CA1 laminar gene expression at rostral, mid–rostrocaudal, and caudal levels in 5xFAD mice (Figs. S7 and S8). Across all three rostrocaudal levels and disease stages examined, male and female mice exhibited highly similar laminar organization, with preservation of overall layer boundaries and subregion-specific spatial patterns.

Given this shared spatial framework, we next examined whether more subtle sex-associated differences emerge at the level of gene expression magnitude at mid-CA1 (HGEA Level 82). Across disease stages and CA1 subregions, male and female 5xFAD mice exhibited largely overlapping laminar gene expression profiles, indicating conserved molecular organization between sexes (Fig. S9). Split-violin analyses revealed highly similar expression distributions for *Lrmp*, *Ndst4*, *Trib2*, and *Peg10* between males and females at early and intermediate disease stages (2mo and 6mo) across all CA1 subregions, motivating subsequent analyses at the advanced disease stage (Fig. S10).

At 14mo, we next assessed whether these sex-associated differences were evident at the level of laminar cell type density. Welch’s t-test with Benjamini–Hochberg FDR correction did not detect statistically significant sex differences in positive-cell density for any CA1 laminar cell types across subregions (Fig. S10). Although some variability was observed across individual animals, these differences did not reach statistical significance after multiple-comparison correction. These results indicate that laminar cell type densities are largely comparable between males and females at advanced stages of disease.

## Discussion

Using a molecularly defined HGEA laminar framework to classify CA1 pyramidal neuron cell types, we show that disease progression in the 5xFAD mouse model does not produce uniform neurodegeneration across the CA1 axis. Instead, distinct laminar populations follow divergent disease trajectories of susceptibility and resilience. Integrating our previously established spatial gene expression mapping approach ^9^ with analyses of cellular abundance and molecular state across disease stage, sex, and rostrocaudal position, this study reveals that disease-associated changes unfold through cell type-specific responses that vary across CA1 subregions. Notably, CA1 laminar architecture remains intact across disease stages despite progressive amyloid deposition, while molecular alterations emerge within specific laminar populations prior to plaque deposition, suggesting early molecular dysregulation associated with 5xFAD mutations. These findings indicate that CA1 pyramidal neuron populations with distinct molecular profiles are differentially affected by APP, PSEN1, and PSEN2 mutations, reframing prior CA1 degeneration phenotypes as cell type-specific responses rather than pyramidal neuron loss due to regional or spatial localization to disease pathology.

### Layer 2 pyramidal neurons exhibit selective vulnerability across disease progression

Prior studies have demonstrated that vulnerability within dorsal CA1 is selective rather than uniform, with preferential degeneration of deep pyramidal neurons reported in tauopathy models ^18^ and superficial pyramidal neurons in epilepsy ^19^. Our findings that CA1 Layer 2 neurons (deep pyramidal neurons in CA1d) are primarily vulnerable in 5xFAD mice suggest convergent vulnerability across amyloid- and tau-associated neuropathology in AD. Although prior studies were limited to dorsal CA1, we found that Layer 2 cell density declines within CA1d, CA1i, and CA1v across disease progression, indicating cell type vulnerability rather than region-specific degeneration.

Amyloid plaque burden in 5xFAD mice becomes enriched in ventral hippocampal regions at mid-to-late stages but is not restricted to specific laminar domains within CA1 ^20^. Our analyses show that Layer 2 cellular and molecular alterations occur across CA1d, CA1i, and CA1v, rather than being restricted to CA1v where plaque burden is greatest. Thus, although Layer 2 vulnerability overlaps spatially with regions of elevated amyloid deposition, selective susceptibility cannot be explained solely by plaque proximity. Similarly, tauopathy models report preferential accumulation of phosphorylated tau within deep CA1d pyramidal neurons compared with superficial populations, followed by degeneration of the deep pyramidal layer ^18^, suggesting that intrinsic properties of CA1 Layer 2 pyramidal neurons confer heightened sensitivity, with pathological substrates amplifying rather than defining susceptibility.

CA1 Layer 2 pyramidal neurons span the entire rostrocaudal axis of CA1 and correspond to the deep pyramidal sublayer in CA1d, but superficial pyramidal layer in CA1i and CA1v ^9^. Most prior studies have focused predominantly on the more rostral part of dorsal CA1 (HGEA CA1d) where ‘superficial’ and ‘deep’ neurons correspond to our CA1 Layers 1 and 2, respectively ^19,21–25^. Consistent with this interpretation, previous studies indicate that deep pyramidal neurons exhibit higher intrinsic firing rates and distinct patterns of network engagement compared with superficial populations ^19,26^. These activity-dependent properties may increase metabolic or synaptic stress, rendering this population particularly sensitive during disease progression.

In addition to physiological features, molecular programs intrinsic to CA1 Layer 2 neurons may contribute to their heightened susceptibility. The Layer 2 marker gene *Ndst4* encodes a bifunctional N-deacetylase/N-sulfotransferase involved in the biosynthesis of heparan sulfate proteoglycans ^27,28^. Unlike the more broadly expressed *Ndst1 and Ndst3*, *Ndst4* exhibits a more restricted expression pattern, as observed in Allen Brain Atlas in situ hybridization data (www.mouse.brain-map.org), suggesting that specific cell types may rely on distinct heparan sulfate modification programs to regulate their signaling environments. Heparan sulfate proteoglycans regulate synaptic interactions and extracellular signaling within neural circuits, influencing how neurons respond to growth factors and other environmental cues ^29^. These molecules are also associated with amyloid-beta deposits and can influence amyloid aggregation and stability ^30,31^. In addition, cell-surface heparan sulfate proteoglycans mediate the binding and uptake of tau aggregates, facilitating their internalization and propagation ^32,33^. Although the specific molecular consequences of reduced *Ndst4* expression in CA1 Layer 2 neurons remain to be determined, these observations suggest that transcriptional alterations in this population may modify extracellular signaling environments that contribute to the selective vulnerability observed during disease progression. Consistent with this interpretation, surviving Layer 2 neurons exhibit reduced *Ndst4* expression, suggesting that the highest-expressing cells are preferentially lost and that the remaining population occupies a diminished molecular state.

### Layer 3 pyramidal neurons exhibit delayed susceptibility during disease progression

CA1 Layer 3 pyramidal neurons display a comparatively modest response profile during disease progression in 5xFAD mice. *Trib2*-positive Layer 3 neurons do not show comparably clear early reductions in positive-cell density or contraction in cellular composition, distinguishing them from the more vulnerable CA1 Layer 2 population. Across CA1i and CA1v, Layer 3 cell density and fractional representation remain relatively preserved at early stages, with a transient increase at 6 mo before declining at later stages. A significant disease-stage effect was detected in CA1v, whereas CA1i exhibited a similar trajectory, but did not reach statistical significance.

At the molecular level, *Trib2* expression similarly shows a comparable pattern, with modest increases at intermediate disease stages followed by significant reductions at later timepoints. This trajectory contrasts with the progressive transcriptional decline observed in Layer 2 neurons and maintained or increasing marker expression observed in Layers 1 and 4. These findings suggest that CA1 Layer 3 pyramidal neurons exhibit delayed susceptibility during disease progression, with changes emerging later and progressing more gradually than those observed in the highly vulnerable Layer 2 population. These observations indicate that CA1 pyramidal neuron responses to amyloid-associated pathology span a continuum rather than a binary pattern of vulnerability versus resilience.

### Relative resilience of Layer 1 and Layer 4 pyramidal neurons across disease progression

In contrast to vulnerable CA1 pyramidal neurons, some pyramidal neuron subtypes exhibit relative preservation during disease progression. Transcriptomic analyses in AD-related models report that superficial CA1 neurons do not show enrichment for dysfunction pathways observed in more vulnerable deep populations ^16^, suggesting potential laminar differences in susceptibility. Within the 5xFAD model, our experiments show CA1 Layer 1 and Layer 4 pyramidal neurons demonstrate relative resiliency and preservation compared to Layer 2 and 3 neurons. *Lrmp*-positive Layer 1 neurons in CA1d and *Peg10*-positive Layer 4 neurons in CA1v and CA1vv comprise an increasing fraction of the surviving pyramidal neuron population across disease progression, in contrast to the decline observed in Layer 2 populations. As vulnerable populations diminish, Layers 1 and 4 represent a larger proportion of the remaining pyramidal neuron pool. This compositional shift reflects relative preservation rather than proliferation. At the molecular level, these populations retain or upregulate marker gene expression within surviving cells, including increased *Lrmp* expression in Layer 1 neurons and *Peg10* expression in Layer 4 neurons. These findings indicate that CA1 pyramidal neuron cell types follow divergent trajectories of vulnerability and resilience which can be distinguished due to the conserved laminar framework of the CA1.

### Rostrocaudal position and sex as covariates of cell type vulnerability

Our findings indicate that the severity of cell type susceptibility in 5xFAD mice varies along the rostrocaudal axis, while sex does not substantially alter the cell type vulnerability hierarchy observed. Along the hippocampal axis, rostral CA1 remains comparatively preserved, whereas mid and caudal regions exhibit progressively greater disruption. Across these regions, Layer 2 pyramidal neurons remain the most severely affected population. This spatial gradient parallels evidence from human studies showing that AD does not affect the hippocampus uniformly along the anterior-posterior axis, with earlier involvement of anterior regions and more extensive degeneration observed in posterior hippocampal domains ^34^.

Biological sex is increasingly recognized as a modifier of AD progression ^35–37^, yet how sex influences vulnerability at the level of defined neuronal cell types remains unclear. Within the framework applied here, laminar organization and cell type densities are highly similar between male and female 5xFAD mice, and quantitative analyses do not reveal significant sex differences across CA1 layers or subregions examined. Together, these findings indicate that while rostrocaudal position influences the magnitude of degeneration, the hierarchy of laminar vulnerability is conserved across sexes and primarily reflects intrinsic cell type identity.

### Comparison to publicly available transcriptomic datasets

External transcriptomic resources were examined to evaluate whether CA1 marker genes were associated with biological programs and functional domains represented in AD datasets. Across multiple AD mouse models in the MODEL-AD resource, CA1 marker genes exhibited consistent enrichment patterns across sexes and disease timepoints, indicating that these genes are reproducibly detected across AD model datasets (Fig. S11a; left). In addition, correlations between 5xFAD molecular signatures and human AD consensus transcriptional clusters linked CA1 marker genes to biological programs associated with extracellular matrix organization, immune processes, neuronal signaling, and cellular stress responses (Fig. S11a; right). Functional domain annotations from the Agora AD Knowledge Portal further supported these associations, connecting CA1 laminar marker genes with conserved biological domains observed in human AD datasets (Fig. S11b).

Consistent with the domain associations observed for *Ndst4* in human AD datasets (Fig. S11b), pathway enrichment analyses were performed on *Ndst4*-high neuronal populations in external human transcriptomic datasets. Cells exceeding the 75th percentile of *Ndst4* expression showed enrichment for pathways related to synaptic transmission, postsynaptic signaling, and glutamatergic receptor activity (Fig. S11c). These findings indicate that *Ndst4*-high neurons are associated with molecular programs linked to neuronal signaling and synaptic function, consistent with the proposed role of *Ndst4*-mediated heparan sulfate modification in regulating extracellular signaling environments within CA1 circuits.

## Methods

### Experimental mouse model

The 5xFAD transgenic mouse model expresses five familial Alzheimer’s disease mutations in human *APP* and *PSEN1* under the neuron-specific Thy-1 promoter, producing rapid amyloid pathology. Experimental cohorts were generated by crossing heterozygous male 5xFAD mice (MMRRC strain #034848-JAX) ^13^ with female C57BL/6 mice (Jackson Labs #000664) generating transgenic 5xFAD offspring and littermate wild-type (WT) controls. Offspring were pair-housed by sex and genotype under standard conditions (21–22° C, 51% humidity, 12 h light/12 h dark cycle, lights on 6:00 a.m.) with ad libitum access to food and water. All procedures were approved by the Institutional Animal Care and Use Committee at the University of Southern California. Sex was incorporated into the study design, and full cohort composition (age, sex, genotype, and n per experiment) is provided in Fig. S1.

### Selection of experimental timepoints

Experimental age timepoints were selected to represent early (2 months), intermediate (6 months), and advanced (14 months) stages of amyloid-associated disease progression in the 5xFAD mouse model, based on established pathological and functional milestones ^14,15^ (AlzForum.org). The 2mo timepoint corresponds to APP upregulation, early amyloid plaque deposition, and gliosis, whereas 6mo represents a transitional stage characterized by robust amyloid accumulation, synaptic marker loss, and early morphological and transcriptional alterations. The 14mo timepoint reflects advanced disease with extensive amyloid pathology and pronounced cellular and molecular disruption. Intervening ages were not included because pathological progression in 5xFAD mice occurs along a continuum without discrete intermediate transitions.

### Probe design and manufacturing

Marker genes were selected based on previously established CA1 laminar gene expression signatures from the Hippocampal Gene Expression Atlas (HGEA) ^9^. The following marker genes were used to label laminar-defined CA1 pyramidal neuron populations: *Lrmp* (Layer 1), *Ndst4* (Layer 2), *Trib2* (Layer 3), and *Peg10* (Layer 4). RNAscope HiPlex v2 probes targeting *Lrmp (*catalog #559391-T8), *Ndst4* (catalog #313631-T6), *Trib2* (catalog #514021-T7), and *Peg10* (catalog #512921-T5) were designed and manufactured by Advanced Cell Diagnostics (ACD) (catalog #324409). Each target probe was assigned a distinct tail corresponding to a cleavable fluorophore channel to enable multiplexed detection within the same tissue section.

### Tissue preparation and RNAscope hybridization

Mice were euthanized with Euthasol and transcardially perfused with fresh 0.9% saline and 4% paraformaldehyde (PFA; pH 9.5). Brains were post-fixed in 4% PFA for 24-48 h at 4° C, embedded in 3% Type 1-B agarose (Sigma-Aldrich), and sectioned into four series of 20 μm-thick coronal sections using a VF-510-0Z Compresstome (Precisionary Instruments). Sections were mounted onto SuperFrost Plus slides (FisherSci, catalog #12-550-15).

In situ hybridization was performed using the RNAscope HiPlex v2 assay according to the manufacturer’s protocol (ACD). Sections were baked at 60° C, dehydrated through graded ethanol (50%, 75%, and 100%), and subjected to target retrieval (RNAscope Target Retrieval Reagent, catalog #322000), followed by protease treatment (Protease III, catalog #322340). Probes were hybridized at 40° C for 2 h, followed by signal amplification (Amp 1-3). Fluorophore hybridization was performed using RNAscope Fluoro T1-T4 v2 reagents (AF488, Dylight 550, Dylight 650, AF750), and sections were counterstained with DAPI. Slides were coverslipped with ProLong Gold antifade mounting medium (FisherSci, catalog #P36930).

### Microscopy imaging and image post-processing

Sections were imaged using an Andor Dragonfly spinning-disk confocal microscope (Oxford Instruments) equipped with a 40x Nikon silicon oil immersion objective. Whole-section tile scans were acquired through the full tissue thickness (15–20 µm) using probe-appropriate fluorescence filter sets. Z-stacks were deconvolved, stitched to generate three-dimensional image volumes, and converted to maximum intensity projections (MIPs) in Fiji for downstream quantitative analysis. Exposure times ranged from 90–120ms, with laser intensities between 5–10%, adjusted according to probe signal.

### Section selection and experimental design

For quantitative analyses, CA1 tissue sections were obtained from male and female mice at 2, 6, and 14 months of age. Cell type density, fraction, and molecular state were quantified at a mid-rostrocaudal CA1 level (HGEA Level 82) containing all four laminar-defined CA1 pyramidal neuron populations. Each animal contributed one representative section and served as the unit of biological replication. Additional rostral and caudal sections were used for visualization of spatial gene expression patterns but were not included in quantitative or statistical analyses.

### Cell segmentation and transcript detection

MIP images were analyzed using QuPath^38^ as previously described ^9^. Regions of interest (ROIs) corresponding to CA1 subregions (CA1d, CA1i, CA1v, CA1vv) were manually annotated to encompass the pyramidal layer. DAPI-labeled nuclei were detected using the QuPath cell detection algorithm, and cellular ROIs were generated by expanding nuclear boundaries to include surrounding cytoplasm (pixel intensity threshold: 2.5 – 5). Subcellular transcript detection was performed using QuPath’s subcellular detection algorithm to identify fluorescent puncta within each gene channel. Detection parameters were adjusted per probe and section to account for signal intensity and background variability. Cell-by-gene count matrices and corresponding cellular ROIs were exported from QuPath for downstream analysis.

### Gene expression normalization and SCAMPR-based analysis

Cell-by-gene detection matrices and corresponding cellular ROI annotations were analyzed in R using the SCAMPR pipeline ^39^. Gene expression values were normalized at the single-cell level by converting area fraction measurements to absolute expression values (Area Fraction x ROI Area / 100) and log-transformed for subsequent analyses. Expression distributions across CA1 subregions, ages, and sexes were visualized using violin plots. Two spatial visualization approaches were generated from the normalized dataset. Topographic maps were constructed by plotting individual cell ROIs according to the medial—lateral and dorsal—ventral coordinates and coloring each cell by the gene with the highest log-normalized expression value; cells lacking detectable expression across markers were unassigned. Spatial distribution maps displayed log-normalized expression of individual genes using a fixed color scale (0—2 log units) to enable comparison across ages.

Pairwise co-expression analyses were performed using log-normalized single-cell expression values aggregated across animals. Adjacent-layer marker pairs (*Lrmp—Ndst4, Ndst4—Trib2, Trib2—Peg10*) were analyzed within CA1 subregions enriched for both markers. Expression values were plotted at the single-cell level and modeled using locally estimated scatterplot smoothing (LOESS) within the SCAMPR co-expression framework. Residuals from the LOESS fit were visualized to identify cells deviating from the modeled relationship. CA1vv was excluded due to enrichment of only one single marker.

### Cell type classification and thresholding

Laminar CA1 pyramidal neuron cell types were classified using gene– and subregion—specific expression thresholds anchored to pooled 2mo WT tissue (n=4), which exhibits stable laminar organization prior to disease-associated changes in gene expression levels. For each marker (*Lrmp, Ndst4, Trib2, Peg10*) and CA1 subregion, thresholds were defined as the median of non-zero normalized expression values measured in pooled 2mo WT tissue and were applied uniformly across all 5xFAD samples and ages. Marker detection was restricted to subregions containing the corresponding laminar population (e.g., *Lrmp* in CA1d; *Ndst4* in CA1d-CA1v; *Trib2* in CA1i–CA1v; *Peg10* in CA1v–CA1vv).

Cell type identity was assigned using an exclusive classification approach: among markers exceeding threshold within a cell, the marker with the highest expression was assigned as the cell type identity, whereas cells not exceeding threshold for any marker were classified as unassigned. Classification quality control metrics were quantified across animals, including gene- and subregion-specific threshold values, assignment distributions, and the proportion of unassigned cells, as well as consistency between assigned cell counts and DAPI-labeled nuclei.

### Cell type abundance and molecular state quantification

Positive-cell density and cell type fractional composition were quantified for each CA1 pyramidal neuron population across CA1 subregions (CA1d, CA1i, CA1v, CA1vv) and ages (2mo, 6mo, 14mo). Positive-cell density was calculated as the number of cells assigned to a given laminar cell type divided by the corresponding ROI area (µm^2^) and expressed as cells/mm^2^. Fractional composition was calculated as the proportion of cells assigned to each laminar cell type relative to the total number of DAPI-segmented nuclei within the same ROI. ROI areas were obtained from QuPath annotation measurements. Molecular state was quantified as the mean log-normalized expression value within cells assigned to each cell type.

### Statistical analysis

Disease-stage differences in positive-cell density, fraction composition, and molecular state were assessed using Kruskal-Wallis tests performed independently for each gene x subregion panel comparing 2mo, 6mo, 14mo 5xFAD mice. P-values were adjusted for multiple comparisons using the Benjamini-Hochberg false discovery rate (FDR) across tested panels within each analysis. Where significant effects were detected, pairwise comparisons were performed using Dunn’s post-hoc test.

Comparisons between age-matched WT and 5xFAD animals were assessed using Wilcoxon rank-sum tests, with p-values corrected using the Benjamini-Hochberg FDR procedure across gene-by-subregion panels. Sex-stratified comparisons within a single disease stage were assessed using Welch’s t-test to account for unequal variances and sample sizes, with p-values corrected using the Benjamini-Hochberg FDR procedure across corresponding gene x subregion panels.

### External transcriptomic dataset analyses

External datasets were examined to evaluate biological programs associated with CA1 cell type marker genes (*Lrmp, Ndst4, Trib2, Peg10*). Marker gene enrichment across AD mouse models was assessed using the MODEL-AD resource (3xTg-AD, 5xFAD, hABKI, Trem2), and correlations between 5xFAD molecular signatures and human AD consensus transcriptional clusters were examined. Functional domain annotations for CA1 marker genes were obtained from the AD Knowledge Portal, which aggregates analyses from human AD datasets.

For pathway enrichment analyses, *Ndst4*-high neurons were defined as cells exceeding the 75th percentile of Ndst4 expression. Gene ontology over-representation analysis (ORA) was performed using the Allen Institute Whole Human Brain (WHB) dataset and gene set enrichment analysis (GSEA) was performed using the Aligning Science Against Parkinson Collaborative Research Network - Post-mortem derived brain sequencing (ASAP-PMDBS) dataset to identify biological pathways associated with *Ndst4*-high neuronal populations.

## Declarations

### Ethics approval and consent to participate

All animal experiment protocols were approved by the USC Institutional Animal Care and Use Committee (IACUC).

### Data availability

The datasets supporting the conclusions of this article are included within the article and its Source data files. Further information is available from the corresponding author upon reasonable request.

### Author contributions

M.P. and M.S.B. designed the experiments. M.P. performed the experiments with the help of A.H. M.P. and S.M. analyzed the data. B.B. and Y.Z. performed analyses of external open-access datasets. M.P. wrote the manuscript draft, M.P. and M.S.B. edited and revised the final manuscript. All authors read and approved the final manuscript.

## Acknowledgements

The authors would like to thank the USC Mark and Mary Stevens Institute for Neuroimaging and Informatics for their guidance and support of the Center of Integrative Connectomics (CIC). The authors also thank members of the Bienkowski Lab for helpful discussions and feedback throughout the course of this study. In addition, the authors acknowledge the publicly available datasets that contributed to this work.

## Supplementary Information

### Supplementary File 1. Supplementary Figures

Contains all supplementary figures and corresponding figure legends.

## References

1. Jack, C. R. et al. Medial temporal atrophy on MRI in normal aging and very mild Alzheimer’s disease. Neurology 49, 786–794 (1997).

2. Yushkevich, P. A. et al. Nearly automatic segmentation of hippocampal subfields in in vivo focal T2-weighted MRI. NeuroImage 53, 1208–1224 (2010).

3. Mueller, S. G. et al. Hippocampal atrophy patterns in mild cognitive impairment and Alzheimer’s disease. Hum. Brain Mapp. 31, 1339–1347 (2010).

4. Dickerson, B. C. et al. Medial temporal lobe function and structure in mild cognitive impairment. Ann. Neurol. 56, 27–35 (2004).

5. Small, S. A., Schobel, S. A., Buxton, R. B., Witter, M. P. & Barnes, C. A. A pathophysiological framework of hippocampal dysfunction in ageing and disease. Nat. Rev. Neurosci. 12, 585–601 (2011).

6. Acosta, D., Powell, F., Zhao, Y. & Raj, A. Regional vulnerability in Alzheimer’s disease: The role of cell-autonomous and transneuronal processes. Alzheimers Dement. 14, 797–810 (2018).

7. Planche, V. et al. Structural progression of Alzheimer’s disease over decades: the MRI staging scheme. Brain Commun. 4, fcac109 (2022).

8. Bartsch, T. et al. Selective Neuronal Vulnerability of Human Hippocampal CA1 Neurons: Lesion Evolution, Temporal Course, and Pattern of Hippocampal Damage in Diffusion-Weighted MR Imaging. J. Cereb. Blood Flow Metab. 35, 1836–1845 (2015).

9. Pachicano, M. et al. Laminar organization of pyramidal neuron cell types defines distinct CA1 hippocampal subregions. Nat. Commun. 16, 10604 (2025).

10. Bienkowski, M. S. et al. Homologous laminar organization of the mouse and human subiculum. Sci. Rep. 11, 3729 (2021).

11. Nielsen, J. V., Blom, J. B., Noraberg, J. & Jensen, N. A. Zbtb20-Induced CA1 Pyramidal Neuron Development and Area Enlargement in the Cerebral Midline Cortex of Mice. Cereb. Cortex 20, 1904–1914 (2010).

12. Gómez-Isla, T. et al. Neuronal loss correlates with but exceeds neurofibrillary tangles in Alzheimer’s disease. Ann. Neurol. 41, 17–24 (1997).

13. Oakley, H. et al. Intraneuronal β-Amyloid Aggregates, Neurodegeneration, and Neuron Loss in Transgenic Mice with Five Familial Alzheimer’s Disease Mutations: Potential Factors in Amyloid Plaque Formation. J. Neurosci. 26, 10129–10140 (2006).

14. Oblak, A. L. et al. Comprehensive Evaluation of the 5XFAD Mouse Model for Preclinical Testing Applications: A MODEL-AD Study. Front. Aging Neurosci. 13, 713726 (2021).

15. Forner, S. et al. Systematic phenotyping and characterization of the 5xFAD mouse model of Alzheimer’s disease. Sci. Data 8, 270 (2021).

16. Alldred, M. J. et al. Hippocampal CA1 Pyramidal Neurons Display Sublayer and Circuitry Dependent Degenerative Expression Profiles in Aged Female Down Syndrome Mice. J. Alzheimers Dis. 100, S341–S362 (2024).

17. Li, H. et al. Loss of SST and PV positive interneurons in the ventral hippocampus results in anxiety-like behavior in 5xFAD mice. Neurobiol. Aging 117, 165–178 (2022).

18. Viney, T. J. et al. Spread of pathological human Tau from neurons to oligodendrocytes and loss of high-firing pyramidal neurons in aging mice. Cell Rep. 41, 111646 (2022).

19. Cid, E. et al. Sublayer- and cell-type-specific neurodegenerative transcriptional trajectories in hippocampal sclerosis. Cell Rep. 35, 109229 (2021).

20. Tsui, K. C. et al. Distribution and inter-regional relationship of amyloid-beta plaque deposition in a 5xFAD mouse model of Alzheimer’s disease. Front. Aging Neurosci. 14, 964336 (2022).

21. Abbaspoor, S. & Hoffman, K. L. Circuit dynamics of superficial and deep CA1 pyramidal cells and inhibitory cells in freely moving macaques. Cell Rep. 43, 114519 (2024).

22. Berndt, M., Trusel, M., Roberts, T. F., Pfeiffer, B. E. & Volk, L. J. Bidirectional synaptic changes in deep and superficial hippocampal neurons following in vivo activity. Neuron 111, 2984–2994.e4 (2023).

23. Cavalieri, D. et al. CA1 pyramidal cell diversity is rooted in the time of neurogenesis. eLife 10, e69270 (2021).

24. Masurkar, A. V. et al. Postsynaptic integrative properties of dorsal CA1 pyramidal neuron subpopulations. J. Neurophysiol. 123, 980–992 (2020).

25. De La Prida, L. M. Potential factors influencing replay across CA1 during sharp-wave ripples. Philos. Trans. R. Soc. B Biol. Sci. 375, 20190236 (2020).

26. Mizuseki, K., Diba, K., Pastalkova, E. & Buzsáki, G. Hippocampal CA1 pyramidal cells form functionally distinct sublayers. Nat. Neurosci. 14, 1174–1181 (2011).

27. Kreuger, J. & Kjellén, L. Heparan Sulfate Biosynthesis: Regulation and Variability. J. Histochem. Cytochem. 60, 898–907 (2012).

28. Aikawa, J. & Esko, J. D. Molecular Cloning and Expression of a Third Member of the Heparan Sulfate/Heparin GlcNAcN-Deacetylase/N-Sulfotransferase Family. J. Biol. Chem. 274, 2690–2695 (1999).

29. Condomitti, G. & De Wit, J. Heparan Sulfate Proteoglycans as Emerging Players in Synaptic Specificity. Front. Mol. Neurosci. 11, 14 (2018).

30. Snow, A. D. & Wight., T. N. PROTEOGLYCANS/GLYCOSAMINOGLYCANS IN THE PATHOGENESIS OF ALZHEIMER’S DISEASE AND OTHER AMYLOIDOSES. Neurobiol. Aging Volume 10, Pages 481-497 (1989).

31. Van Horssen, J., Wesseling, P., Van Den Heuvel, L. P., De Waal, R. M. & Verbeek, M. M. Heparan sulphate proteoglycans in Alzheimer’s disease and amyloid-related disorders. Lancet Neurol. 2, 482–492 (2003).

32. Holmes, B. B. et al. Heparan sulfate proteoglycans mediate internalization and propagation of specific proteopathic seeds. Proc. Natl. Acad. Sci. 110, (2013).

33. Stopschinski, B. E. et al. Specific glycosaminoglycan chain length and sulfation patterns are required for cell uptake of tau versus α-synuclein and β-amyloid aggregates. J. Biol. Chem. 293, 10826–10840 (2018).

34. Hrybouski, S. et al. Aging and Alzheimer’s disease have dissociable effects on local and regional medial temporal lobe connectivity. Brain Commun. Volume 5, (2023).

35. Aggarwal, N. T. & Mielke, M. M. Sex Differences in Alzheimer’s Disease. Neurol. Clin. 41, 343–358 (2023).

36. Sil, A. et al. Sex Differences in Behavior and Molecular Pathology in the 5XFAD Model. J. Alzheimers Dis. 85, 755–778 (2022).

37. Sundermann, E. E. et al. Sex differences in Alzheimer’s-related Tau biomarkers and a mediating effect of testosterone. Biol. Sex Differ. 11, 33 (2020).

38. Bankhead, P. et al. QuPath: Open source software for digital pathology image analysis. Sci. Rep. 7, 16878 (2017).

39. Ali Marandi Ghoddousi, R., Magalong, V. M., Kamitakahara, A. K. & Levitt, P. SCAMPR, a single-cell automated multiplex pipeline for RNA quantification and spatial mapping. Cell Rep. Methods 2, 100316 (2022).

