## Supplementary File for "Differential vulnerability of CA1 pyramidal neuron cell types in the 5xFAD Alzheimer’s disease mouse model"

### Supplementary File 1. Supplementary Figures

| Sample sizes by age, sex, and anteroposterior level (HGEA) | | | | | |
|---|---|---|---|---|---|
| **Age** | **HGEA Level** | **Anteroposterior Position** | **Male (n)** | **Female (n)** | **Total (n)** |
| **HGEA Level 82*** | | | | | |
| 2 mo | HGEA Level 82* | Mid CA1 | 2 | 3 | 5 |
| 6 mo | HGEA Level 82* | Mid CA1 | 3 | 3 | 6 |
| 14 mo | HGEA Level 82* | Mid CA1 | 4 | 5 | 9 |
| **Total** | **HGEA Level 82*** | **Mid CA1** | **9** | **11** | **20** |
| **HGEA Level 72** | | | | | |
| 2 mo | HGEA Level 72 | Rostral CA1 | 1 | 1 | 2 |
| 6 mo | HGEA Level 72 | Rostral CA1 | 1 | 1 | 2 |
| 14 mo | HGEA Level 72 | Rostral CA1 | 1 | 1 | 2 |
| **Total** | **HGEA Level 72** | **Rostral CA1** | **3** | **3** | **6** |
| **HGEA Level 89** | | | | | |
| 2 mo | HGEA Level 89 | Caudal CA1 | 1 | 1 | 2 |
| 6 mo | HGEA Level 89 | Caudal CA1 | 1 | 1 | 2 |
| 14 mo | HGEA Level 89 | Caudal CA1 | 1 | 1 | 2 |
| **Total** | **HGEA Level 89** | **Caudal CA1** | **3** | **3** | **6** |
| **All levels** | | | | | |
| **Grand total** | **All levels** | | **15** | **17** | **32** |

*HGEA Level 82 represents the primary anteroposterior level used for all quantitative (statistical) analyses.*

**Fig. S1. Sample sizes by age, sex, and anteroposterior level (HGEA)**
Sample sizes for both 5xFAD and WT mice analyzed across three experimental timepoints and stratified by sex and Hippocampal Gene Expression Atlas (HGEA) level. The 5xFAD cohort includes animals analyzed at three disease-ages (2mo, 6mo, and 14mo). HGEA Level 82 corresponds to mid-rostrocaudal CA1 and represents the primary anteroposterior level used for all quantitative (statistical) analyses in this study. HGEA Levels 72 (rostral CA1) and 89 (caudal CA1) are included for anatomical and spatial context. Values indicate the number of animals per group (male, female, and total).

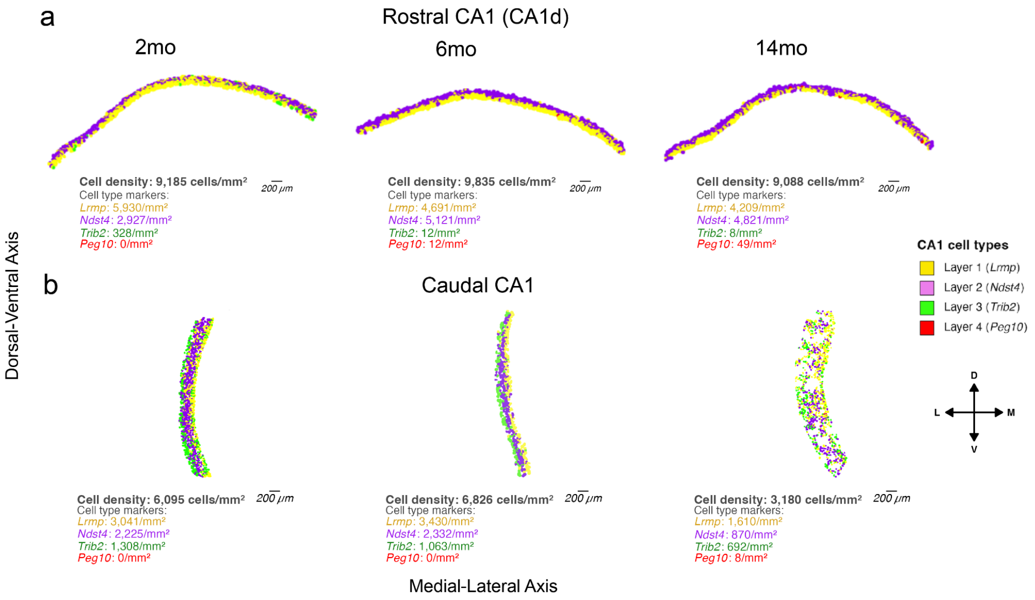


**Fig. S2. Laminar gene expression layers across the CA1 longitudinal axis during disease progression in 5xFAD mice.**

Representative topographic maps illustrating the spatial organization of laminar gene expression layers across the CA1 longitudinal axis in male 5xFAD mice. Rostral CA1d (HGEA Level 72; **a**) and caudal CA1 (HGEA Level 89; **b**) are shown at 2mo, 6mo, and 14mo. Individual cells are plotted according to their medial–lateral (x-axis) and dorsal–ventral (y-axis) anatomical positions within the CA1 pyramidal layer and are colored according to the gene with the highest log-normalized expression value (L*rmp, Ndst4, Trib2*, or *Peg10*), corresponding to laminar cell type populations. Cells lacking detectable expression across markers are unassigned. Total cell density and marker gene-associated cell densities (cells/$mm^2$) are indicated below each map. Scale bars, 200μm. Source data are provided with this paper as supplementary files.

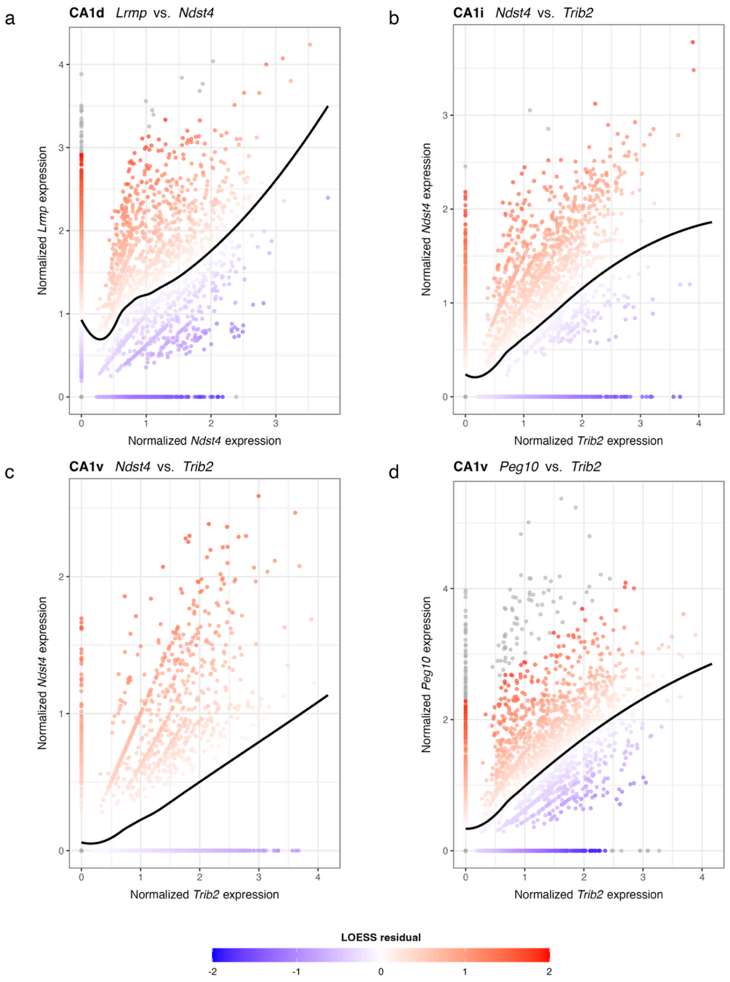


**Fig. S3. Gene coexpression relationships among CA1 subregions at 14mo in 5xFAD mice.**
Scatter plots showing pairwise relationships between log-normalized gene expression values for laminar marker genes within CA1 subregions at 14mo in 5xFAD mice. Each point represents a single pyramidal neuron, plotted using normalized expression values for the indicated gene pair. Quantification is based on HGEA Level 82 (mid-CA1) sections from 14mo 5xFAD mice (n=9; 4 male, 5 female), with all detected pyramidal neurons included in the analysis. Panels show gene pairs corresponding to adjacent marker-associated layers within specific CA1 subregions: (**a**) CA1d, *Lrmp* vs. *Ndst4*; (**b**) CA1i, *Ndst4* vs. *Trib2*; (**c**), CA1v, *Ndst4* vs. *Trib2*; and (**d**) CA1v, *Peg10* vs. *Trib2*. A locally estimated scatterplot smoothing (LOESS) curve (black line) summarizes the non-linear relationship between expression values for each gene pair. Points are colored by LOESS residuals, indicating deviation from the fitted trend (blue, lower than expected; red, higher than expected expression relative to the fit). Source data are provided with this paper as supplementary files.

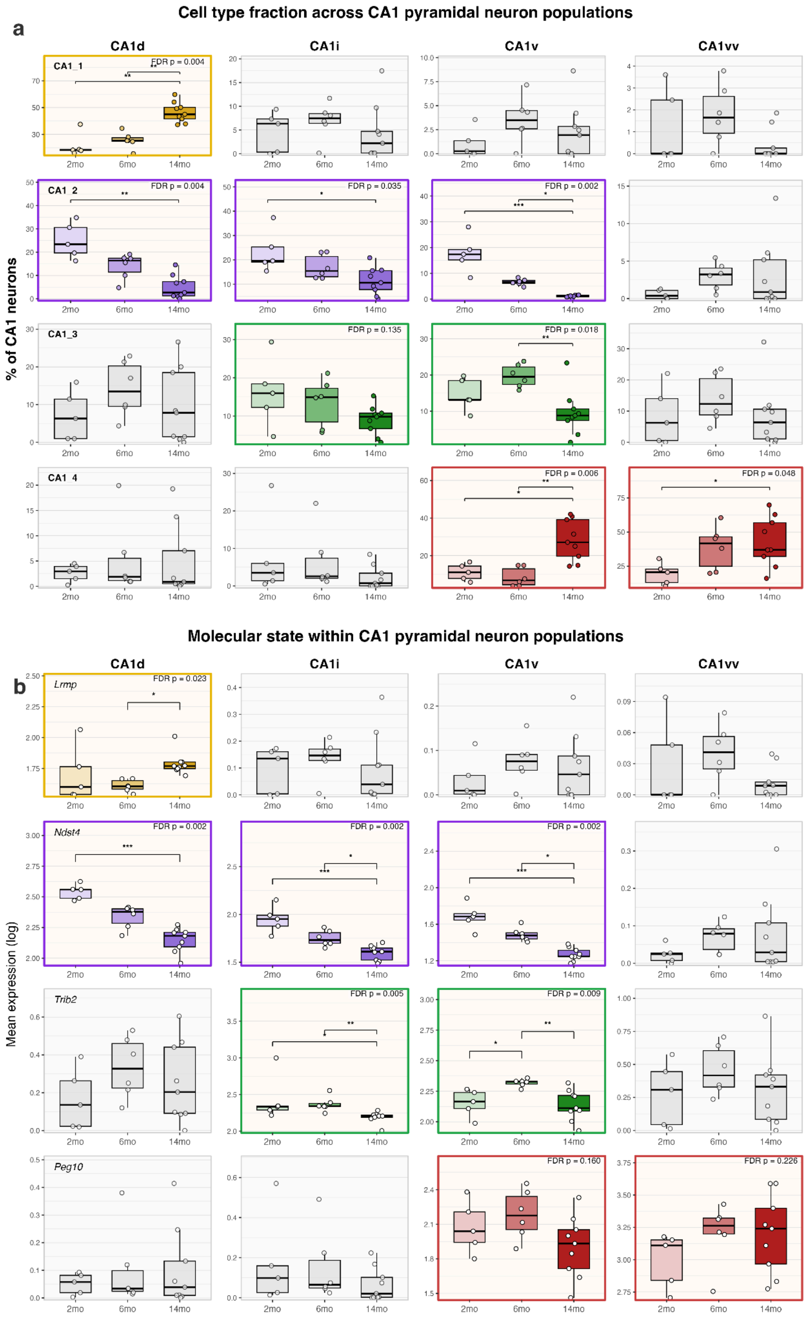


**Fig. S4. Cell type fraction and molecular state across CA1 pyramidal neuron populations during disease progression in 5xFAD mice.**

**a,** Fractions of CA1 pyramidal neuron cell types across disease-associated ages (with the following sample sizes: 2mo, n=5 (2 male, 3 female); 6mo, n=6 (3 male, 3 female); 14mo, n=9 (4 male, 5 female)) in 5xFAD mice, expressed as the percentage of total pyramidal neurons within each CA1 subregion (CA1d, CA1i, CA1v, CA1vv). Cell types are defined according to laminar classification (Layers 1-4) using gene-specific expression thresholds described in the Methods. **b,** Molecular state within each CA1 pyramidal neuron cell type, quantified as mean log-normalized expression of the corresponding marker gene within each classified cell type. Boxplots summarize per-animal values, with individual points representing single animals. Statistical comparisons across ages were performed using Kruskal-Wallis tests comparing 2mo, 6mo, and 14mo 5xFAD mice within each panel, with Benjamini-Hochberg FDR correction applied across all panels. Where significant effects were detected, pairwise comparisons were performed using Dunn's test. FDR-adjusted p-values are shown, and significant pairwise comparisons are indicated by brackets and asterisks. Exact p-values and pairwise Dunn post-hoc comparisons are provided in the corresponding supplementary file.

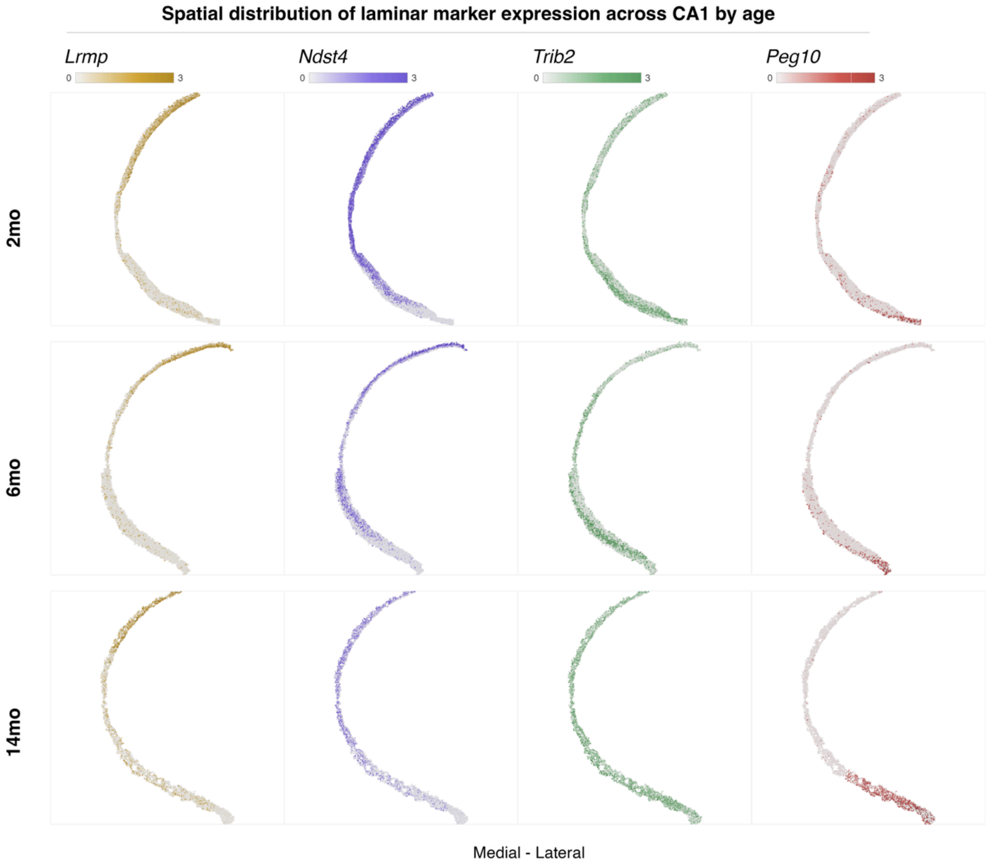


**Fig. S5. Spatial gene expression patterns across CA1 by age in 5xFAD mice.**
Spatial maps showing medial-lateral distributions of log-normalized gene expression of *Lrmp*, *Ndst4*, *Trib2*, and *Peg10* across three disease-associated timepoints (2mo, 6mo, and 14mo) in 5xFAD mice. Each column corresponds to a gene and each row to an age group. Expression values are plotted at the single-cell level, with color intensity reflecting relative gene expression along the CA1 medial-lateral axis. Source data are provided with this paper as supplementary files.

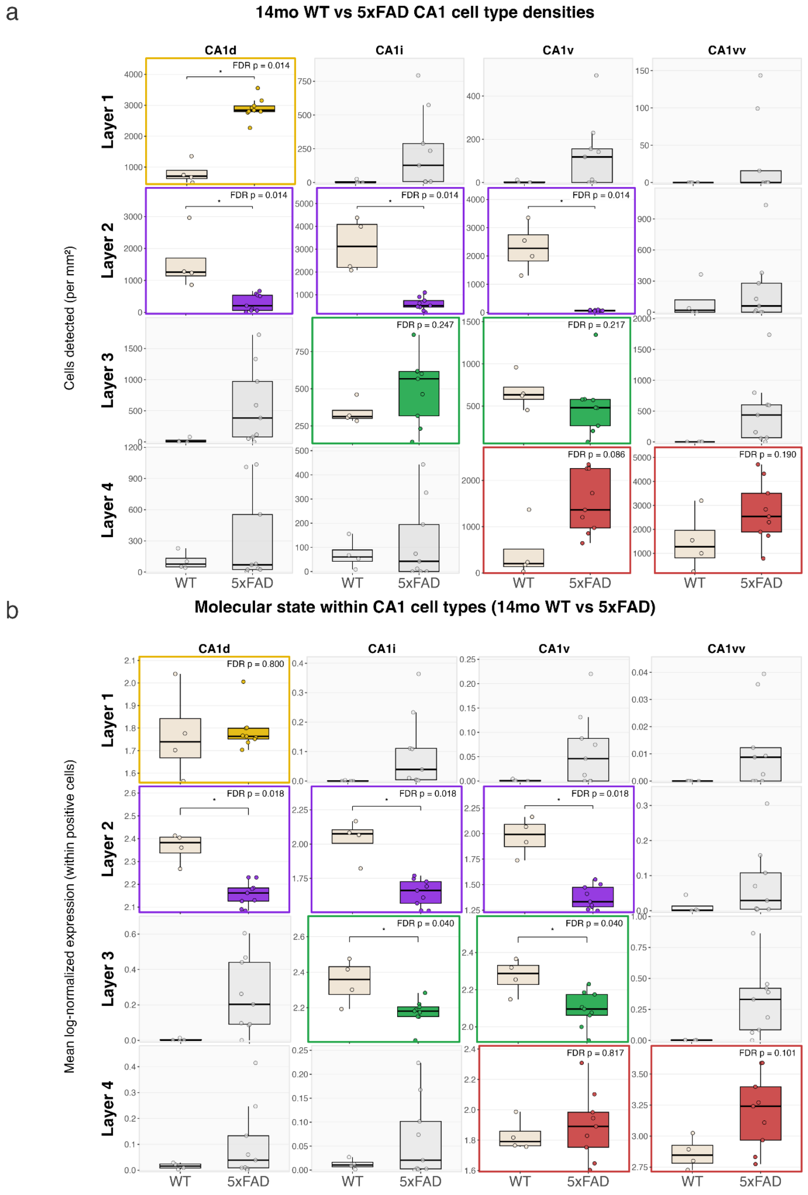


**Fig. S6. Disease-associated changes in CA1 pyramidal neuron cell types at advanced age.**
Boxplots summarize per-animal measurements with individual points representing single animals. Age-matched littermate WT and 5xFAD mice were compared at 14mo using the CA1 laminar cell type classification framework defined using WT reference thresholds (see Methods) (5xFAD group: n=9; 4 male, 5 female; WT group: n=4; 1 male, 3 female). Quantification is based on HGEA Level 82 (mid-CA1) sections. **a**, Positive-cell density (cells/mm$^2$) for each CA1 pyramidal neuron cell type across subregions. **b**, Molecular state within classified cells, defined as the mean log-normalized expression of the defining marker within each cell type. Rows correspond to laminar CA1 cell types (Layers 1-4) and columns correspond to CA1 subregions (CA1d, CA1i, CA1v, CA1vv). Cell type–subregion combinations corresponding to expected laminar gene expression layers are highlighted with colored borders and gene-tinted fills, whereas non-corresponding cell type–subregion combinations are shown in grayscale. Statistical significance was assessed using Wilcoxon rank-sum tests comparing WT and 5xFAD mice within each corresponding cell type–subregion panel, with Benjamini-Hochberg FDR correction applied across tested panels. FDR-adjusted p-values are shown, and significant pairwise comparisons are indicated by brackets and asterisks. Source data are provided with this paper as supplementary files.

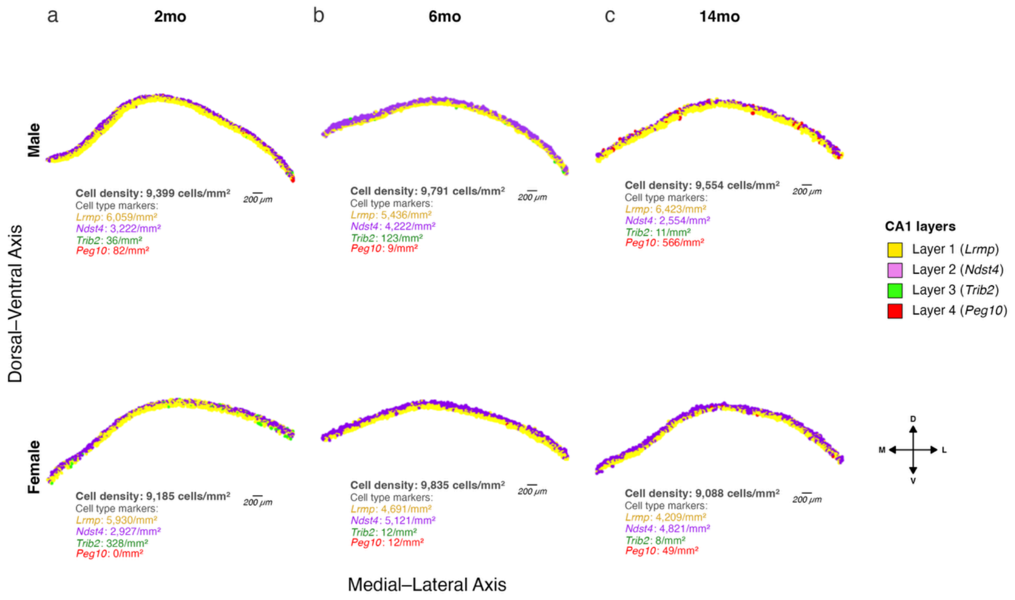


**Fig. S7. Rostral CA1 laminar gene expression layers across disease progression in 5xFAD mice.** Representative topographic maps illustrating the spatial organization of laminar gene expression layers in rostral CA1 (HGEA Level 72) from male (top row) and female (bottom row) 5xFAD mice at 2mo. **a**, 6mo **b**, and **c,** 14mo. Individual cells are plotted according to their medial—lateral (x-axis) and dorsal—ventral (y-axis) anatomical positions within the CA1 pyramidal layer and are colored according to the gene with the highest log-normalized expression value (*Lrmp*, *Ndst4*, *Trib2*, or *Peg10*), corresponding to laminar cell type populations. Cells lacking detectable expression across markers are unassigned. Total cell density and marker gene-associated cell densities (cells/mm$^2$) are indicated below each map. Scale bars, 200 μm. Source data are provided with this paper as supplementary files.

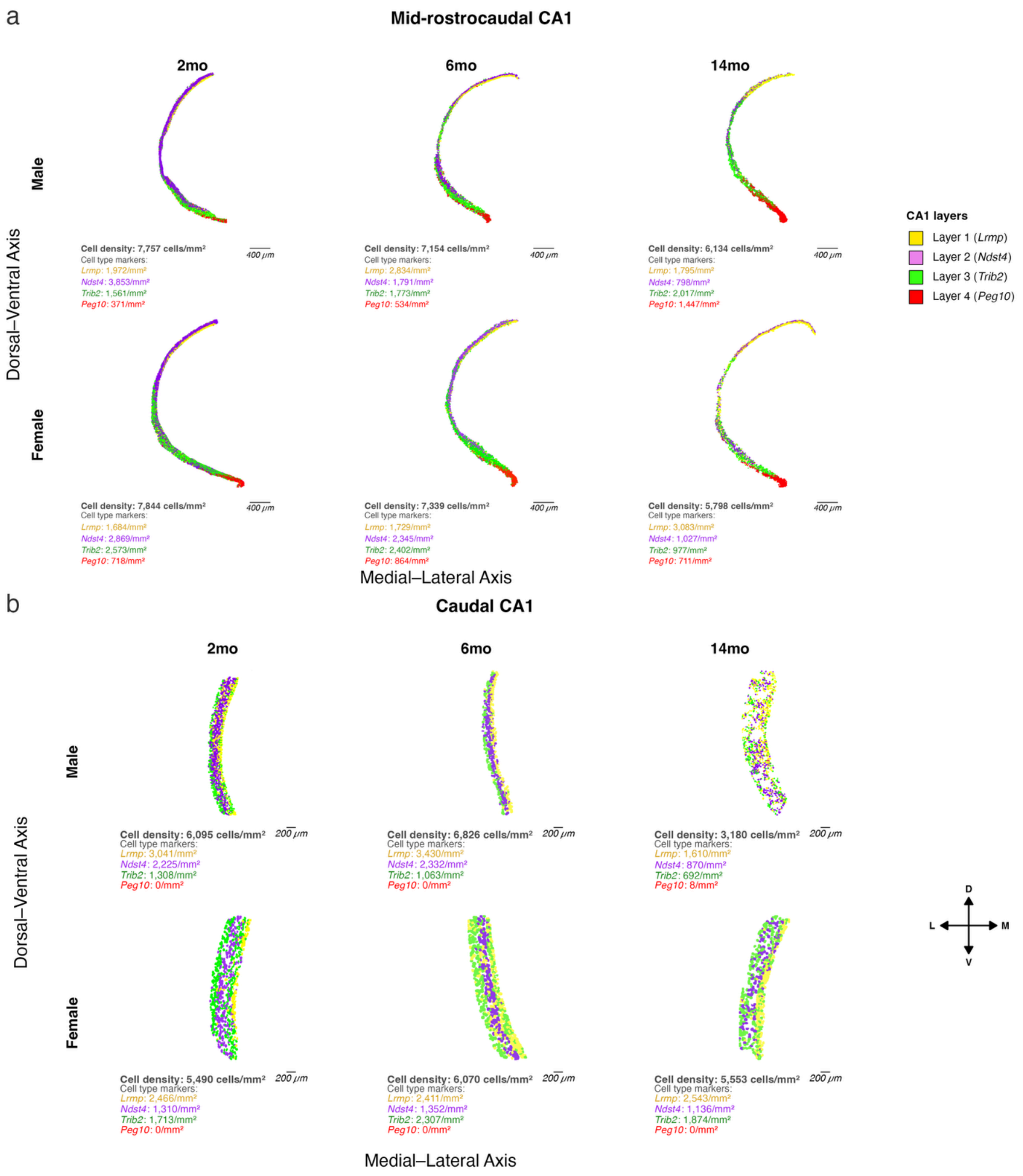


**Fig. S8. Mid-rostrocaudal and caudal CA1 laminar gene expression layers across disease progression in 5xFAD mice.**
Representative topographic maps illustrating the spatial organization of laminar gene expression layers in mid-rostrocaudal CA1 (HGEA Level 82; **a**) and caudal CA1 (HGEA Level 89; **b**) from male (top rows) and female (bottom rows) 5xFAD mice at 2mo, 6mo, and 14mo. Individual cells are plotted according to their medial—lateral (x-axis) and dorsal—ventral (y-axis) anatomical positions within the CA1 pyramidal layer and are colored according to the gene with the highest log-normalized expression value (*Lrmp*, *Ndst4*, *Trib2*, or *Peg10*), corresponding to laminar cell type populations. Cells lacking detectable expression across markers are unassigned. Total cell density and marker gene-associated cell densities (cells/mm$^2$) are indicated below each map. Scale bars, 400 μm (**a**) and 200 μm (**b**). Source data are provided with this paper as supplementary files.

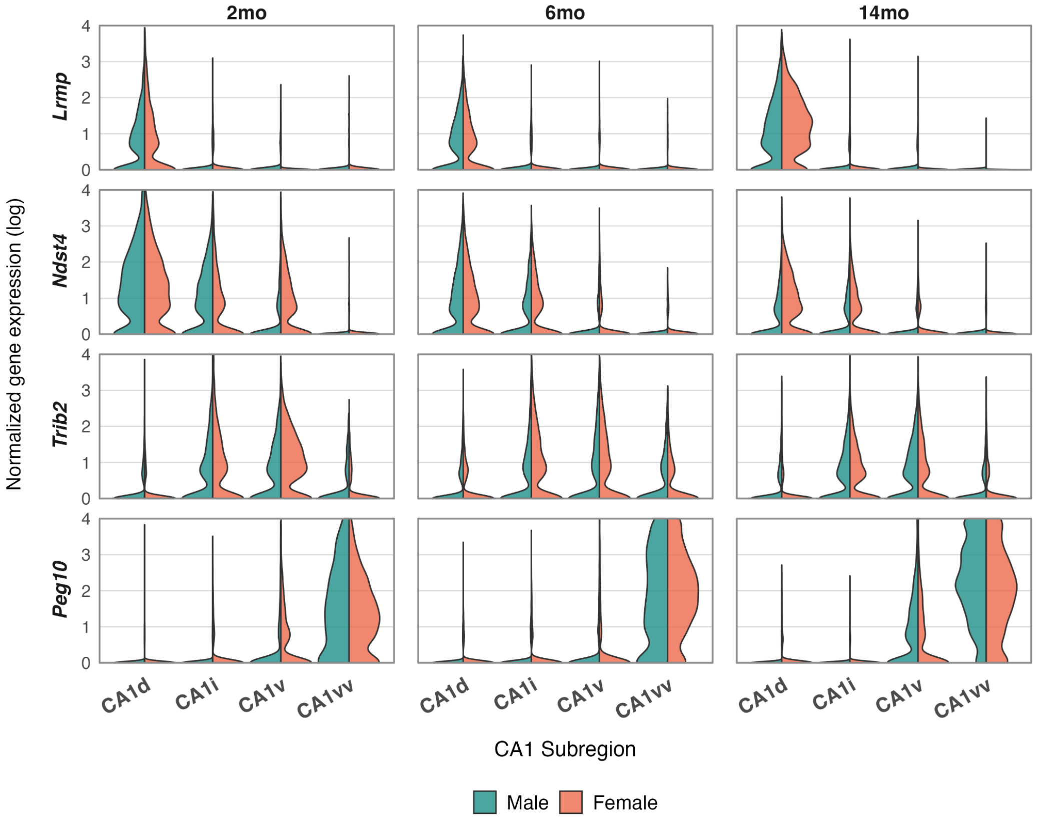


**Fig. S9. Sex differences in CA1 gene expression profiles across subregions during disease progression in 5xFAD mice.**

Split violin plots showing log-normalized gene expression levels of *Lrmp*, *Ndst4*, *Trib2*, and *Peg10* in male and female 5xFAD mice across CA1 subregions (CA1d, CA1i, CA1v, and CA1vv) at 2mo, 6mo, and 14mo. Rows correspond to individual genes and columns correspond to age groups. Within each CA1 subregion, expression distributions for male (teal) and female (salmon) mice are shown side-by-side. Quantification is based on the following sample sizes: 2mo, n=5 (2 male, 3 female); 6mo, n=6 (3 male, 3 female); 14mo, n=9 (4 male, 5 female). Source data are provided with this paper as supplementary files.

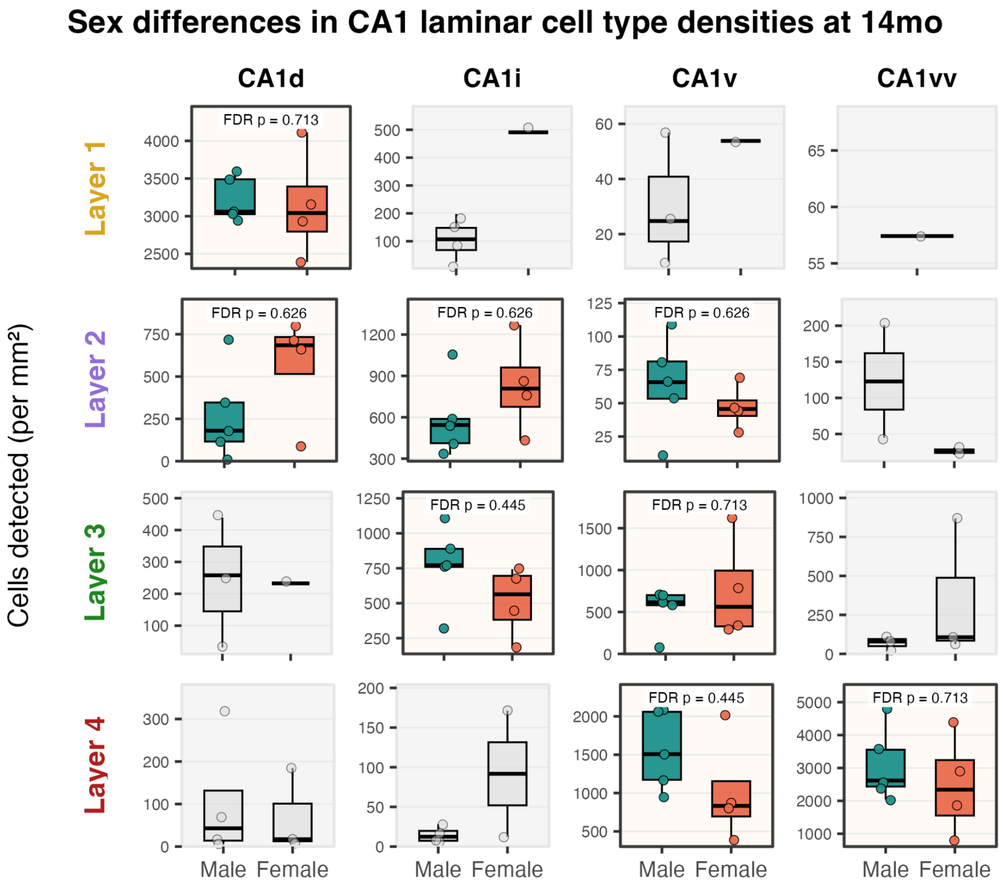


**Fig. S10. Sex differences in CA1 cell type density at 14mo in 5xFAD mice.**
**a,** Sex-stratified positive-cell densities of CA1 pyramidal neuron cell types across CA1 subregions (CA1d, CA1i, CA1v, CA1vv) in 14mo 5xFAD mice. Boxplots summarize per-animal positive-cell densities (cells/mm$^2$), with individual points representing single animals. Quantification is based on HGEA Level 82 (mid-CA1) sections with the following sample sizes: males, n=4; females n=5. Laminar cell types correspond to Layer 1 (*Lrmp*), Layer 2 (*Ndst4*), Layer 3 (*Trib2*), and Layer 4 (*Peg10*). Layer-subregion combinations consistent with expected laminar organization are highlighted with colored borders, whereas non-corresponding combinations are shown in grayscale. Statistical significance was assessed using two-tailed Welch's t-test comparing males and females within each corresponding cell type–subregion panel, with Benjamini-Hochberg FDR correction applied across tested panels. FDR-adjusted p-values are shown. Source data are provided with this paper as supplementary files.

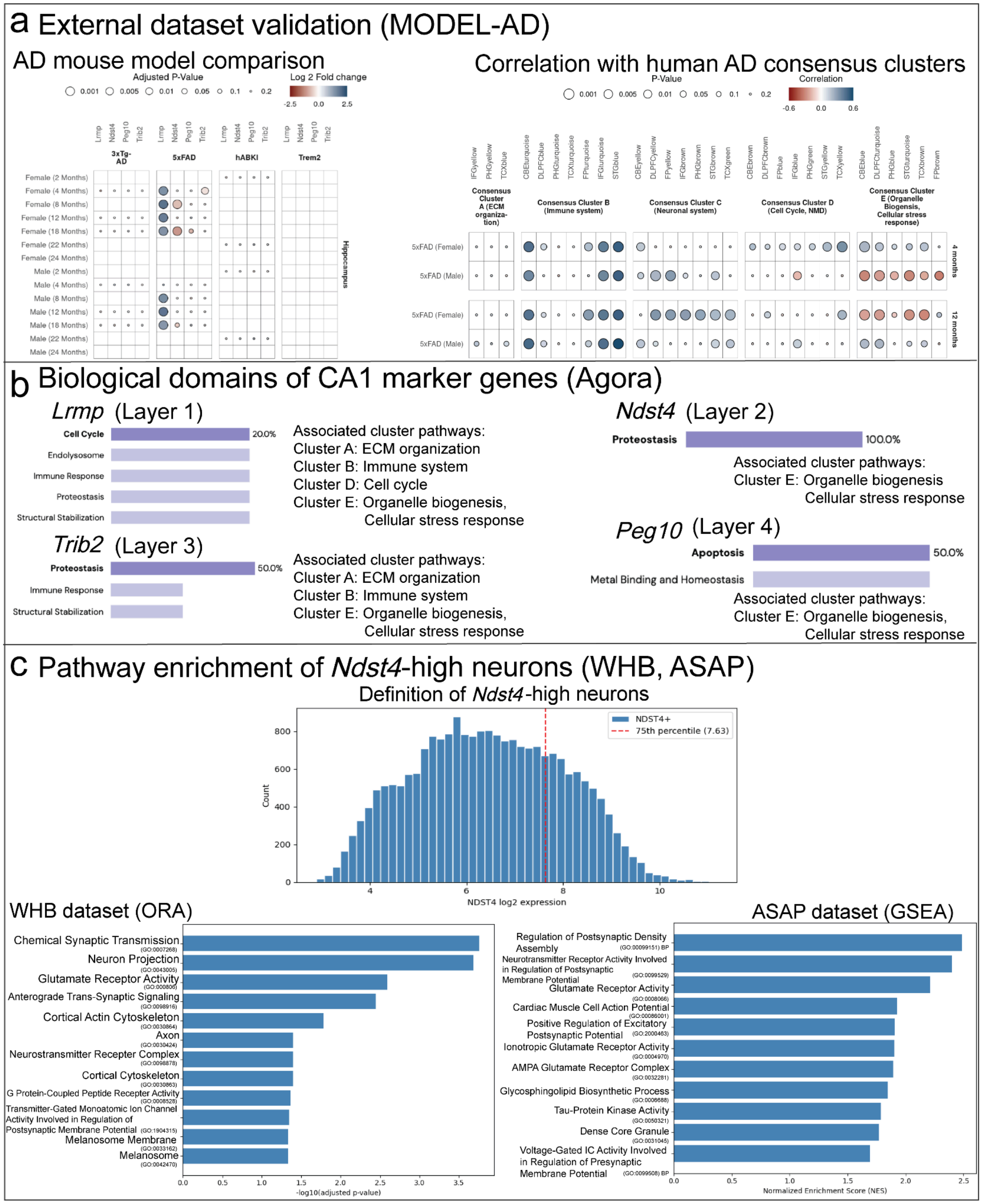


**Fig. S11. External datasets support CA1 marker gene associations and biological pathways.**
**a,** External dataset validation (MODEL-AD). Left, enrichment of CA1 marker genes (*Lrmp, Ndst4, Trib2, Peg10*) across AD mouse models from the MODEL-AD resource (3xTg-AD, 5xFAD, hABKI, Trem2). Bubble size indicates adjusted p-value and color indicates log fold change. Right, correlation of 5xFAD molecular signatures with human AD consensus transcriptional clusters. **b,** Biological domains associated with CA1 marker genes in human AD datasets. Functional domain annotations for CA1 marker genes obtained from the Agora AD Knowledge Portal. **c,** Pathway enrichment of *Ndst4*-high neurons in human datasets. *Ndst4*-high cells were defined using the 75$^{th}$ percentile expression threshold. Pathway enrichment was assessed using gene ontology over-representation analysis (ORA) in the WHB dataset and gene set enrichment analysis (GSEA) in the ASAP-PMDBS dataset. GO term identifiers are displayed in smaller font for readability.
